# Aging-associated Alterations in the Gene Regulatory Network Landscape Associate with Risk, Prognosis and Response to Therapy in Lung Adenocarcinoma

**DOI:** 10.1101/2024.07.02.601689

**Authors:** Enakshi Saha, Marouen Ben Guebila, Viola Fanfani, Katherine H. Shutta, Dawn L. DeMeo, John Quackenbush, Camila M. Lopes-Ramos

**Affiliations:** Department of Biostatistics, Harvard T. H. Chan School of Public Health, Boston, MA 02115, USA; Channing Division of Network Medicine, Brigham and Women’s Hospital, Boston, MA, USA 02115; Department of Medicine, Harvard Medical School, Boston, MA 02115, USA; Department of Data Science, Dana-Farber Cancer Institute, Boston, MA 02115, USA

**Author notes:** **Corresponding Author:** Camila M Lopes-Ramos.

## Abstract

Aging is the primary risk factor for many individual cancer types, including lung adenocarcinoma (LUAD). To understand how aging-related alterations in the regulation of key cellular processes might affect LUAD risk and survival outcomes, we built individual (person)-specific gene regulatory networks integrating gene expression, transcription factor protein-protein interaction, and sequence motif data, using PANDA/LIONESS algorithms, for both non-cancerous lung tissue samples from the Genotype Tissue Expression (GTEx) project and LUAD samples from The Cancer Genome Atlas (TCGA). In GTEx, we found that pathways involved in cell proliferation and immune response are increasingly targeted by regulatory transcription factors with age; these aging-associated alterations are accelerated by tobacco smoking and resemble oncogenic shifts in the regulatory landscape observed in LUAD and suggests that dysregulation of aging pathways might be associated with an increased risk of LUAD. Comparing normal adjacent samples from individuals with LUAD with healthy lung tissue samples from those without LUAD, we found that aging-associated genes show greater aging-biased targeting patterns in younger individuals with LUAD compared to their healthy counterparts of similar age, a pattern suggestive of age acceleration. This implies that an accelerated aging process may be responsible for tumor incidence in younger individuals. Using drug repurposing tool CLUEreg, we found small molecule drugs with potential geroprotective effects that may alter the accelerating aging profiles we found. We also observed that, in contrast to chronological age, a network-informed aging signature was associated with survival and response to chemotherapy in LUAD.

**Graphical abstract:** 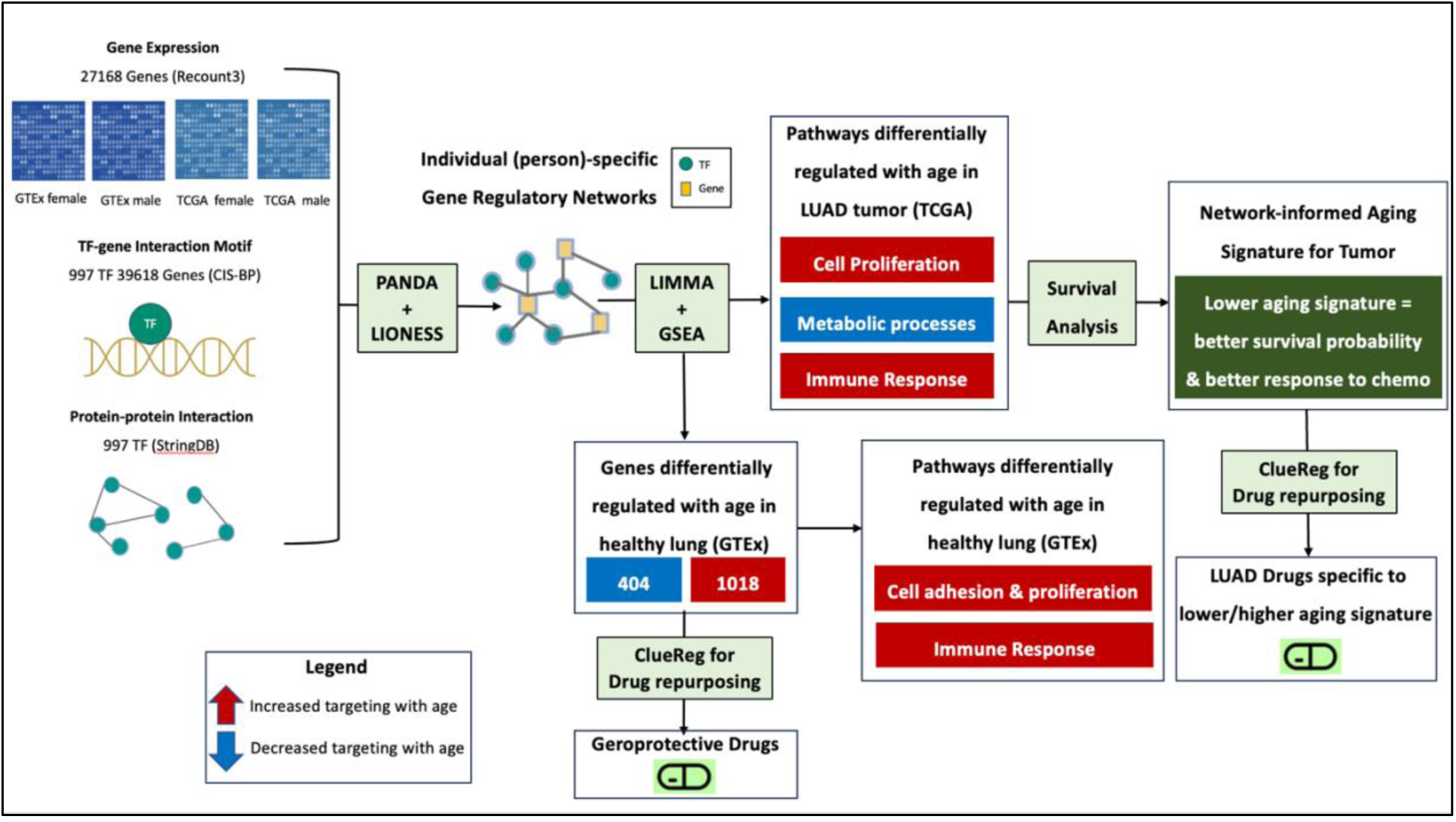

## Introduction

Lung cancer is second only to breast cancer worldwide in annual incidence and is the leading cause of cancer death. Lung cancer risk increases with age and as the average age of the population increases worldwide, the prevalence of lung cancer is expected to continue growing [1]. In 2021, 75% of lung cancer fatalities were reported in individuals aged 65 and older [2]. While lung adenocarcinoma (LUAD) in younger adults is often diagnosed at more advanced stages compared to those in older adults [3], elderly individuals have more comorbidities and tend to be less tolerant of certain cancer therapeutics than younger individuals [4]. These differences are likely the result of aging-induced alterations in the regulation of key cellular processes [5], but the mechanism by which age shifts the gene regulatory landscape to alter lung cancer risk and survival outcome is largely unknown. In this paper we address this critical gap in our understanding by building individual (person)-specific gene regulatory networks to gain insights into aging related changes in gene regulation that might influence the risk and prognosis of LUAD across all age groups. Additionally, we explore how aging-associated regulatory changes are further accelerated by tobacco smoking history, since lung diseases including LUAD are more prevalent among individuals with a history of smoking, compared to individuals who have never smoked in their lifetime [6].

As transcription factors (TF) have been established as known drivers of aging [7], to understand aging-associated heterogeneity in the gene regulatory landscape of LUAD tumors, we identified biological pathways that are differentially regulated by TFs in tumors from individuals of different ages and investigated if any of these age-associated changes in regulatory networks might influence survival and the response to chemotherapy, potentially favoring individuals exhibiting regulatory signatures akin to those found in younger individuals. We further validated our findings in two independent datasets of non-cancerous lung tissue and LUAD tumors.

Most studies investigating the role of aging in lung adenocarcinoma have focused on the mutational landscape of tumor among individuals across various age groups [8, 9]. Tumor mutations in several genes, including *CDKN2A, KRAS*, *MDM2*, *MET*, and *PIK3CA,* have been found to increase in frequency with the individual’s age, while the frequencies of mutation in *ALK*, *ROS1*, *RET* and *ERBB2* show a decreasing trend with age [10]*. ALK* and *EGFR* mutations are high among younger individuals with LUAD, especially among females and nonsmokers [11, 12, 13]. Analysis of somatic interactions has indicated that EGFR-positive samples in younger individuals are more prone to concurrent mutations in PIK3CA, MET, TP53, and RB1 when compared to older individuals [10]. Age may influence both the number of mutations in a tumor and their evolutionary timing [14]. While germline mutations are more commonly identified in tumors from younger individuals, tumors in older individuals appear to be predominantly influenced by somatic mutations [15]. Such mutations clearly play a role in cancer risk and prognosis, acting in part, by altering the activity of biological pathways associated with cancer. However, changes in these pathways can only be partially explained by known mutations, indicating that other mechanisms of pathway activation might play a significant role [16].

Despite some studies in lung cancer that have reported altered expression of genes linked to survival [17, 18], as per our knowledge, there has not been any research investigating aging-associated alterations in gene regulatory networks that influence the risk and prognosis of LUAD and lung cancer in general. We addressed this gap in understanding by using the network-modeling approaches, PANDA [19] and LIONESS [20], to derive person-specific gene regulatory networks for non-cancerous lung tissue samples from the Genotype Tissue Expression project (GTEx) and LUAD tumor samples from The Cancer Genome Atlas (TCGA), with a focus on evaluating age-associated genes and their regulation by TFs. This approach was motivated by multiple earlier network-modeling analyses that identified disease relevant regulatory features in both healthy tissues as well as in tumor [21, 22, 23].

By analyzing individual-specific regulatory networks, we found increased TF targeting of pathways related to intracellular adhesion, cell proliferation, and immune response with age in healthy lung tissue. These aging-associated alterations are further increased by tobacco smoking and resemble oncogenic shifts in the regulatory landscape observed in LUAD tumors, thereby suggesting a potential association between aging-associated dysregulation of biological pathways and an elevated risk of developing LUAD.

Using a web-based drug repurposing tool CLUEreg, we also found potential geroprotective small molecule drug candidates that may be useful in reducing the risk of LUAD by reversing the aging-associated regulatory signatures. By constructing a network-informed aging signature for tumor samples based on the TF-targeting patterns of key biological pathways significantly changing with age in LUAD tumors, we found that a lower aging signature is associated with better survival probability and higher chemotherapy efficacy. In contrast, chronological age was not predictive of survival, thus demonstrating that the aging signature captures aspects of tumor biology not captured by chronological age alone. Using CLUEreg, we also found distinct small molecule drug candidates tailored to LUAD samples with varying aging signatures. In conclusion, our findings not only highlight the mechanisms underlying increased risk and poorer prognosis of LUAD associated with aging-induced gene regulatory alterations, but also establish a potential avenue for leveraging individual-specific gene regulatory networks in designing personalized therapeutic interventions.

## Results

### Identifying Aging-associated Gene Regulatory Alterations in Healthy Human Lung and Geropropective Drug Candidates

We inferred gene regulatory networks linking TFs to target genes using gene expression data from non-cancerous lung tissue samples from GTEx. By analyzing these networks, we identified several genes (**Figure 2**) and biological pathways (**Figure 3**) that are differentially targeted by TFs as a function of age, in the lung. Among these genes, there are 1018 that exhibit significantly increased targeting by TFs with age (p-value <0.05). Most significant among them are *NNAT* [24], *FBLN7* [25], *SH3BP1* [26], *CNTN1* [27], *THEM5* [28], and *FOXP4* [29]; upregulation of these genes has been previously reported to be associated with cell proliferation and poorer prognosis in multiple cancers. We also find 404 genes that exhibit significantly decreased targeting by TFs with age (p-value <0.05) including *DUSP15* [30], *ALDH1L2* [31], *HPD* [32], *GSTT2* [33], *FOXI3* [34], and *ZIC2* [35], all of which have been shown to be influential in predicting tumor progression and therapeutic efficacy in various cancer types, including non-small cell lung cancer.

**Figure 1:**
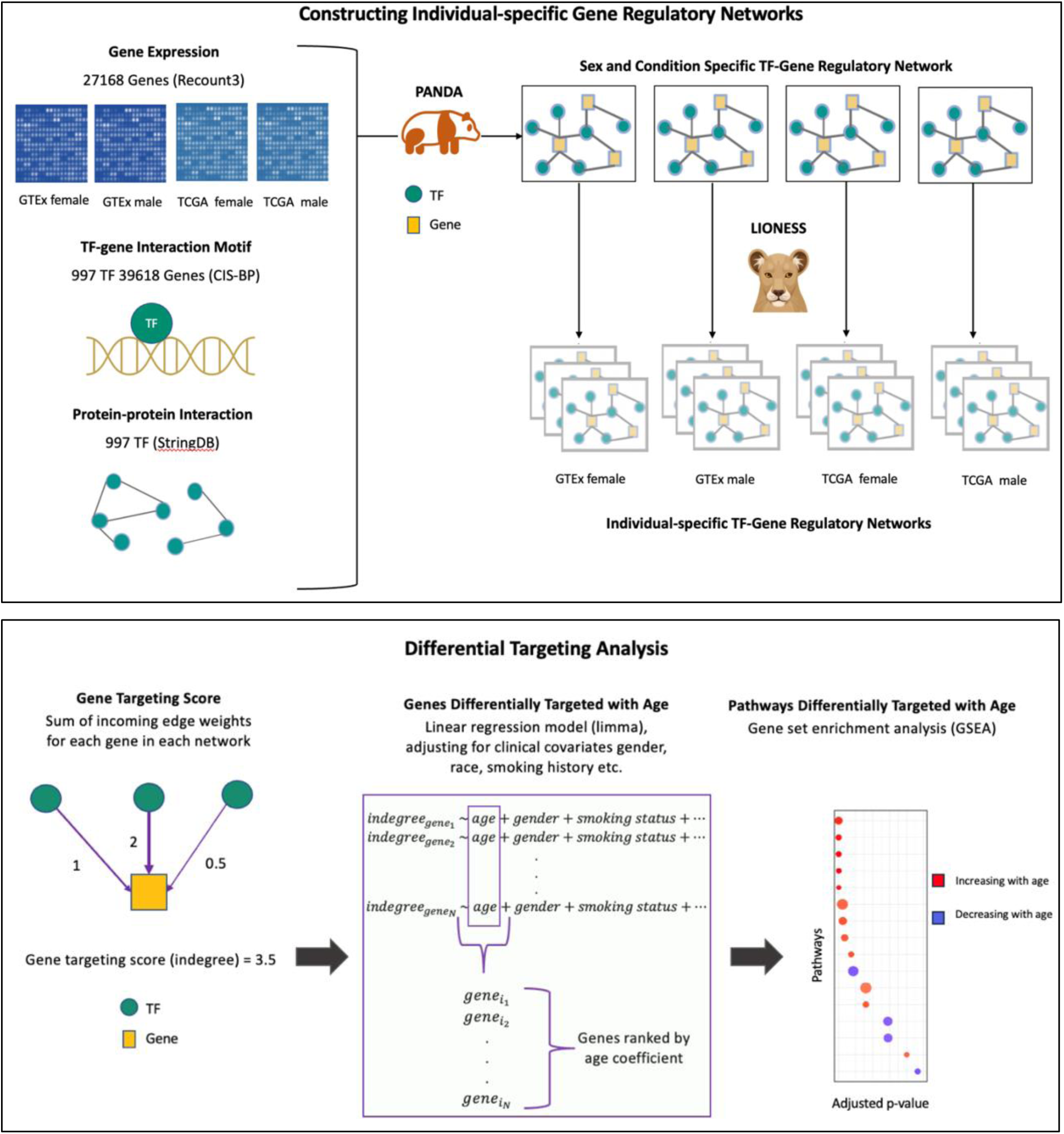
Schematic overview of the study. Top box, overview of our approach to constructing individual specific networks using PANDA and LIONESS which integrate information on protein-protein interactions (PPIs) between transcription factors (TFs), prior information on TF-Gene motif binding, and gene expression data – in this case, from GTEx lung tissues and TCGA LUAD primary tumor samples - downloaded from Recount3. Bottom box, overview of the differential targeting analysis.

**Figure 2:**
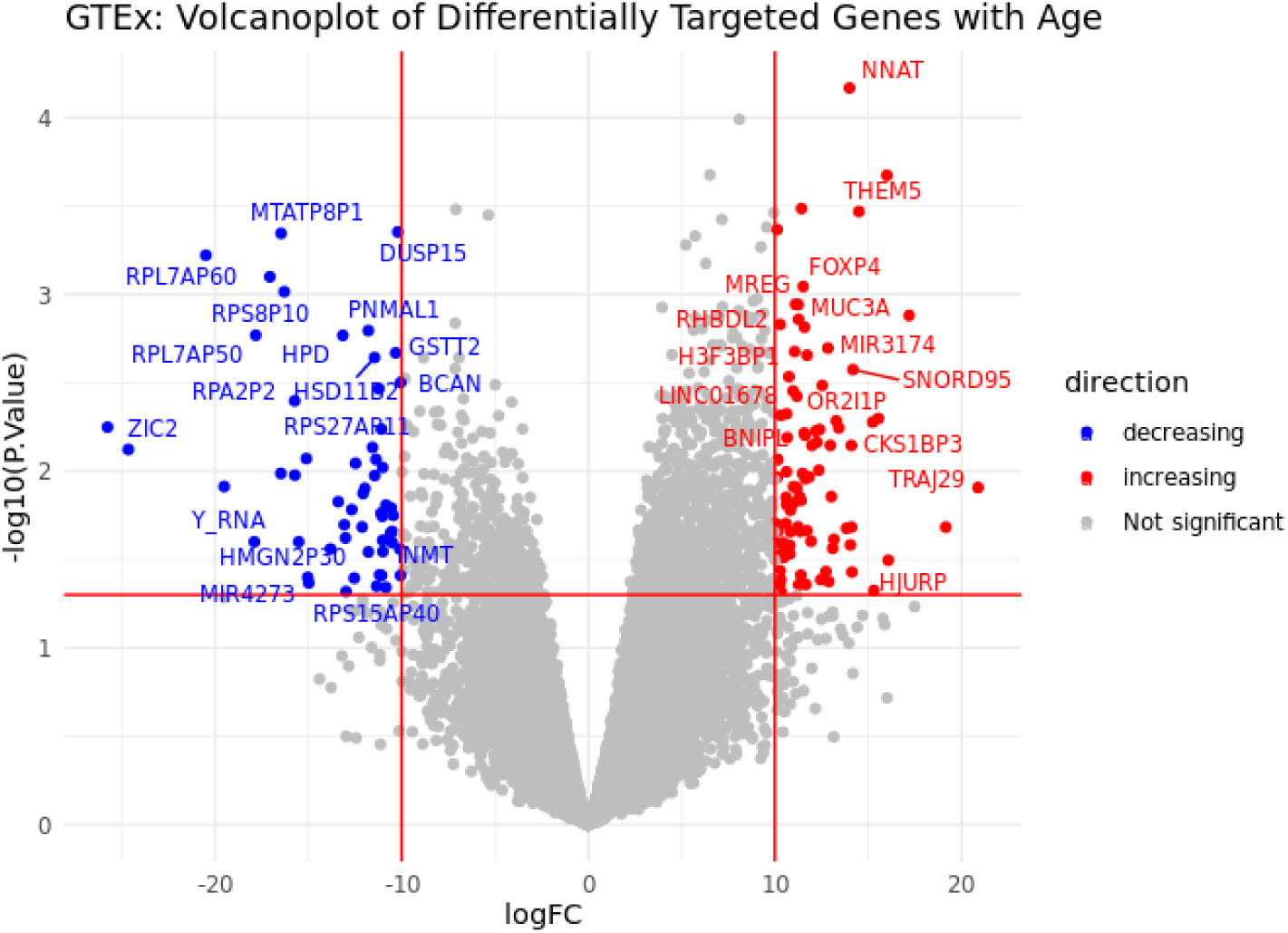
Volcano plot of genes that are differentially (increasingly or decreasingly) targeted by TFs over varying age in lung tissue samples from GTEx. The x-axis represents log fold change (logFC), which is defined as the change in gene indegree in response to a unit change in age. The y-axis represents negative of logarithm of p-values (-log10(P.Value)).

**Figure 3:**
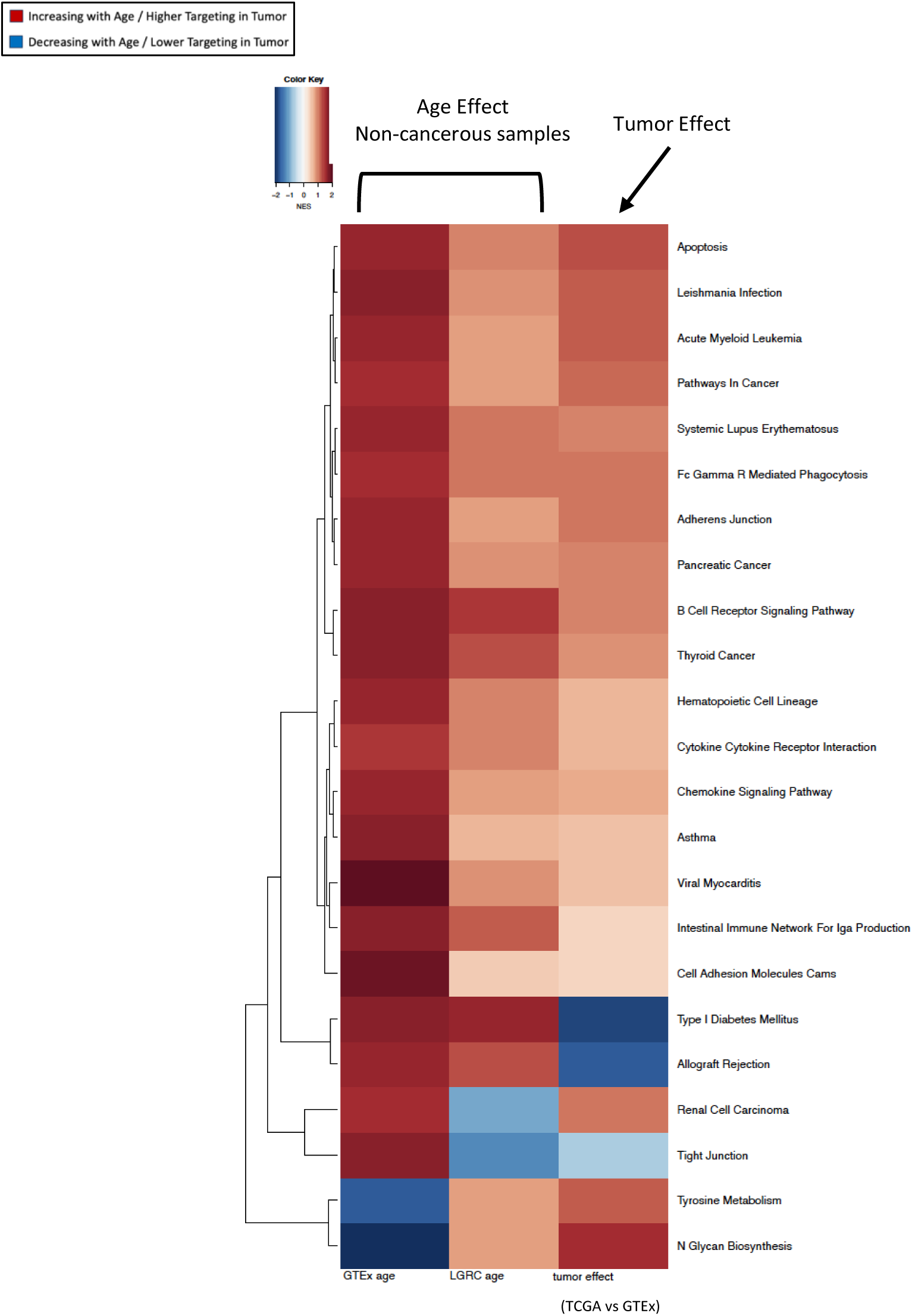
Heatmap of normalized enrichment scores (NES) for pathways that are significantly (at FDR cutoff 0.05) differentially targeted by transcription factors with age among non-cancerous lung samples (GTEx). The first two columns exhibit NES from GSEA based on the age coefficients from the limma analysis of GTEx and LGRC. The third column shows NES from GSEA based on difference between tumor samples from TCGA and healthy samples from GTEx.

We performed gene set enrichment analysis (GSEA) with genes, ranked by how much their targeting patterns change with age, and found (**Figure 3** leftmost column) that biological pathways associated with intracellular adhesion and cell proliferation, cell growth, and death have increasing TF targeting with age, including pathways annotated to adherens junction, apoptosis, hematopoietic cell lineage, cell adhesion molecules, and pathways in cancer. Pathways associated with immune response, including B-cell receptor signaling pathway, cytokine-cytokine receptor interaction, chemokine signaling pathway and intestinal immune network for IgA production, also show increased TF targeting with age. We confirmed these findings on an independent dataset LGRC (**Figure 3** middle column).

Using genes that are differentially targeted by age as input to a web-based drug repurposing tool CLUEreg [36], we identified 150 small molecule drug candidates (Supplementary Material S2) with potential to reverse the aging-associated regulatory alterations in the gene regulatory networks from non-cancerous lung samples. Some of these drugs, henceforth referred to as “geroprotective drugs”, including Carnosol [37], Curcumin [38], Cucurbitacin B [39], Isonicotinamide [40], Meclofenoxate [41], Scriptaid [42], and Withaferin A [43] have already been shown to have potential geroprotective effects in various animal models, including humans. Among these 150 geroprotective drug candidates, we found several FDA-approved anti-cancer drugs, including Trametinib, Doxorubicin, Alisertib, Actinomycin-d, Toremifene, and Plumbagin, as well as several investigational drugs with potential anti-tumor effects including Avrainvillamide analogs [44], aurora kinase inhibitors (MK-5108, AT-9283) [45], Avicins [46], HMN-214 [47], Chaetocin [48], ron kinase inhibitors [49], and Linifanib [50], among others.

### Tobacco Smoking is Associated with Accelerated Aging

To explore whether tobacco smoking is associated with an acceleration of the aging process, we compared the gene regulatory networks from non-cancerous samples, between individuals with a history of smoking and individuals who have never smoked in their lifetime. We split the 1422 aging-associated genes we had previously identified into two sets: genes that exhibit increasing TF targeting with age (1018 genes) and those that show decreasing TF targeting (404 genes). For every gene in these two sets, we used limma [51], to compute the age coefficient (**Figure 4**) in a linear model for individuals with and without a history of smoking.

**Figure 4:**
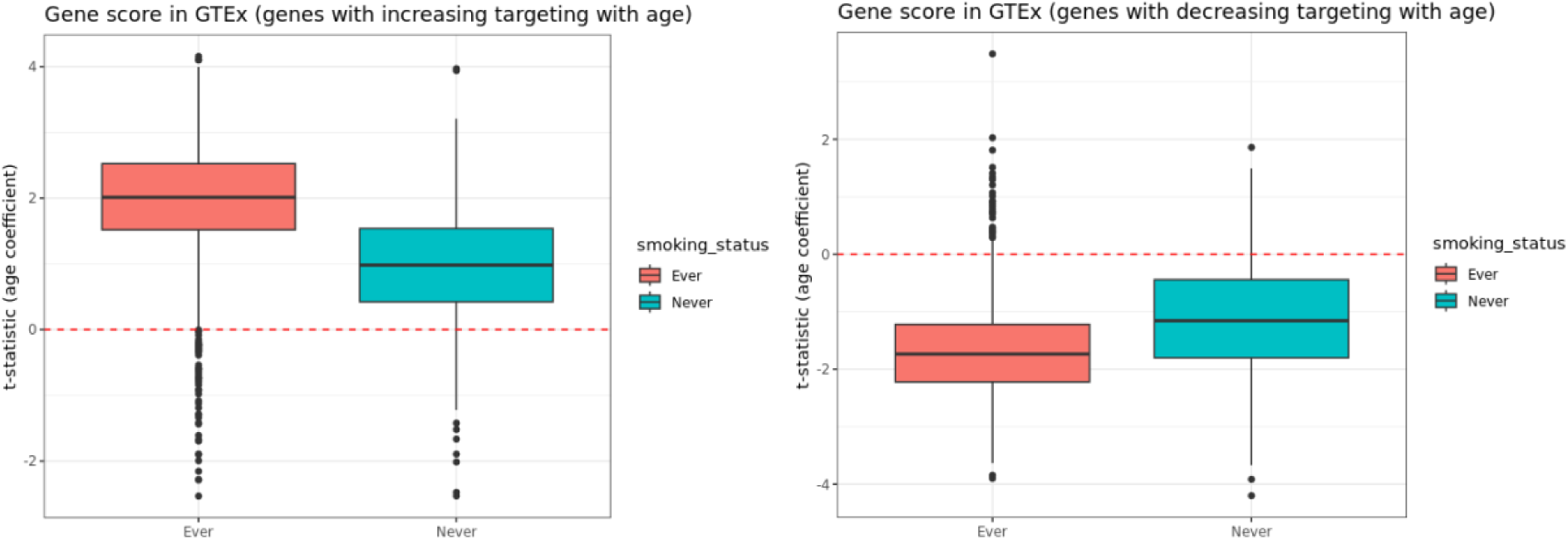
Boxplot of the rates of change in TF-targeting with age in GTEx (designated by the t-statistics from the limma analysis with interaction between age and smoking status) (*Left*) for 1018 genes that are increasingly targeted with age in healthy human lung (based on evidence from GTEx) and (*Right*) the same boxplot for 404 genes that are decreasingly targeted with age in healthy human lung (based on evidence from GTEx).

We found that for genes with increasing age-associated TF targeting, the t-statistics of the age coefficients among individuals with a history of smoking have significantly greater positive values than among never-smokers (p-value < 2.2e-16). Similarly, for genes with decreasing TF-targeting with age, the t-statistic of the age coefficients among individuals with a history of smoking have significantly (p-value < 2.2e-16) larger negative values than those among individuals who have never smoked. In other words, for both kinds of aging-associated genes, the age gradients are significantly steeper for the individuals with a history of smoking, than those for the individuals who have never smoked in their lifetimes. The steeper age gradients mean that the rate of change in gene regulation with age is faster among individuals with a history of smoking, comparted to individuals who have never smoked.

To visually represent the continuous changes in gene regulation with age, we plotted the aging trajectories (as described in Methods) for the non-cancerous samples from GTEx (**Figure C.1** in the Supplementary Material). For the genes which are increasingly targeted by TFs with age (left plot on **Figure C.1**), the slope of the regression line is steeper among individuals who have a history of smoking, compared to individuals who have never smoked. Although for genes that are decreasingly targeted by TFs with age (right plot of **Figure C.1**), we do not observe any significant difference between the slopes of the aging trajectories among individuals with or without a history of smoking. Nevertheless, these findings indicate that tobacco smoking is associated with an acceleration of the aging-induced alterations in gene regulation. We also validated these findings in the non-cancerous lung samples from the independent LGRC (a.k.a. GSE47460) validation dataset (**Figure C.2**).

Taken together, we find that even in individuals without evidence of lung cancer, there are aging-associated changes in the TF-targeting of genes that are further accelerated by tobacco smoking, which may be linked to an increased risk of developing LUAD at a younger age among individuals with a history of smoking.

### Aging-associated Gene Regulatory Alterations in Non-cancerous Lung Resemble Oncogenic Gene Regulatory Shifts Observed in LUAD

To understand how aging-related changes in the topology of gene regulatory networks might be linked to an increased risk of LUAD, we analyzed age-associated changes in network density around the neighborhoods of proto-oncogenes and tumor suppressor genes (downloaded from the COSMIC database [52]). We used linear modeling (implemented in limma) and calculated the t-statistics of the age coefficients (normalized age gradients) for the list of oncogenes and tumor suppressor genes (TSG) (**Figure 5**) and found that among the healthy samples in GTEx, TF-targeting increases with age on average for both oncogenes and TSGs (Wilcoxon rank sum test gives a p-value of 3.908e-09 for oncogenes and 0.001164 for tumor suppressor genes).

**Figure 5:**
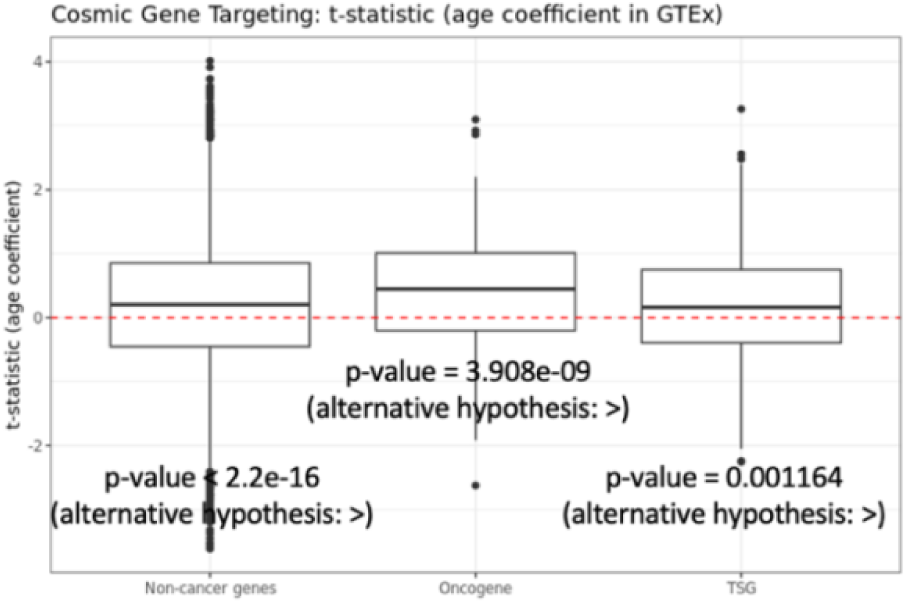
Boxplot of the t-statistics of the age coefficient associated with the oncogenes and tumor suppressor genes (listed in the COSMIC database) from the limma analysis in GTEx. Positive value means these genes are targeted more with age on average and negative value means these genes are targeted less with age on average. For comparison, we also show the same t-statistics for non-cancer genes (genes that are not annotated as oncogenes and/or tumor suppressor genes in the COSMIC database). Each boxplot ranges from the upper and lower quartiles with the median as the horizontal line. Outliers are marked by points. In our analysis we include genes that are explicitly marked as either “Oncogene” or “TSG” respectively in the COSMIC database, thus excluding all the genes that can work either as an oncogene or as a tumor suppressor gene, depending on the mutation. The reported p-values correspond to the hypothesis testing with respect to alternative hypotheses reported in parentheses. The alternative hypotheses “<” or “>” denote the hypotheses “mean > 0” and “mean < 0” respectively.

For comparison, it should be noted that, non-cancer genes (that is, genes not annotated as oncogene or TSG in the COSMIC database) are also increasingly targeted by TFs with age (p-value of Wilcoxon rank sum test is 2.2e-16), meaning that the gene regulatory networks inferred for individuals in GTEx, increase in TF regulatory density as the age of the individual increases. Nevertheless, the mean TF-targeting with aging is the greatest for oncogenes (aging slope of oncogenes is significantly larger than those of non-cancer genes and tumor suppressor genes with p-values being equal to 0.001271 and <2.2e-16 respectively). This indicates that although changes in regulation are a natural consequence of aging, the greatest changes occur in the regulatory neighborhoods of oncogenes, including genes that are common drug targets [53] in LUAD, such as *MYCN*, *ERBB3*, and *AKT1* (**Figure C.3**).

To explore the association of aging with LUAD risk, we compared the targeting patterns of aging-associated biological pathways between non-cancerous samples from GTEx and LUAD tumor samples from TCGA. We observe that aging-associated pathways involved in cell adhesion, cell proliferation and immune response (except for type I diabetes mellitus and allograft rejection pathways), are also highly targeted in LUAD tumor, compared to non-cancerous lung (**Figure 3** rightmost column). This indicates that increased TF-targeting of these pathways with age might be a contributing factor to an elevated risk of developing LUAD among older adults and that those with the greatest regulatory targeting of these pathways might be at the greatest risk.

LUAD among younger individuals, although less frequent, is often detected at more advanced stages compared to their older counterparts [3]. Given that we found age-acceleration of gene targeting to correlate with LUAD, we tested whether LUAD in younger individuals was also associated with patterns of accelerated aging, compared to healthier individuals of similar age. To confirm this hypothesis, we compared the TF-targeting pattern of 1422 aging-associated genes in normal adjacent lung samples from individuals with LUAD (from TCGA) versus non-cancerous lung samples from GTEx (**Figure 6**). For the 1018 genes that exhibited increased targeting with age, we found that their mean TF-targeting was significantly higher (p-value <2.2e-16) in the normal adjacent lung samples from younger individuals (age less than median age 66) with LUAD, compared to individuals of similar age without LUAD (**Figure 6** left). In contrast, for older individuals with LUAD (age greater than median age 66) we did not find a significantly higher mean targeting of aging genes compared to healthy individuals of similar age.

**Figure 6:**
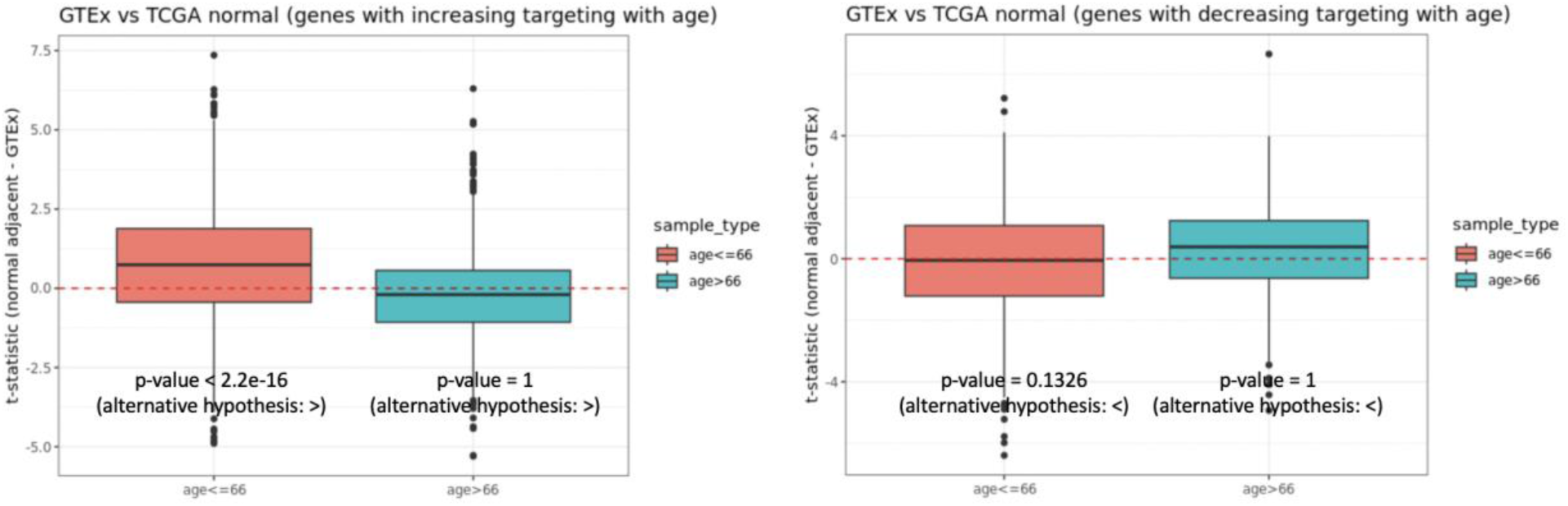
Boxplot of the difference in TF-targeting of genes in TCGA normal adjacent samples compared to GTEx non-cancerous lung for (*Left*) 1018 genes that are increasingly targeted with age in healthy human lung (based on evidence from GTEx) and (*Right*) for 404 genes that are decreasingly targeted with age in healthy human lung (based on evidence from GTEx). The reported p-values correspond to the hypothesis testing with respect to alternative hypotheses reported in parentheses. The alternative hypotheses “<” or “>” denote the hypotheses “mean > 0” and “mean < 0” respectively.

In other words, the gene regulatory patterns observed in the normal-adjacent lung tissues of younger individuals with LUAD are more like those found in older individuals, than they are to their healthy counterparts of the same age. This suggests that LUAD in younger individuals may be driven, in part, by age-accelerated changes in gene regulation, and that this acceleration may also be associated with more aggressive tumor biology at diagnosis we see in younger individuals.

### Biological Pathways Differentially Regulated in LUAD Tumor across Varying Age

We analyzed gene regulatory networks of LUAD samples from TCGA and performed GSEA to identify biological pathways that are differentially regulated by TFs with age (**Figure 7** left column). We found several pathways involved in cell signaling and cell proliferation that were increasingly targeted by TFs with age. When we compared these differentially targeted pathways to those differentially targeted with age in non-cancerous lung samples from GTEx, we found that both had identified the pathway associated with cell adhesion molecules.

**Figure 7:**
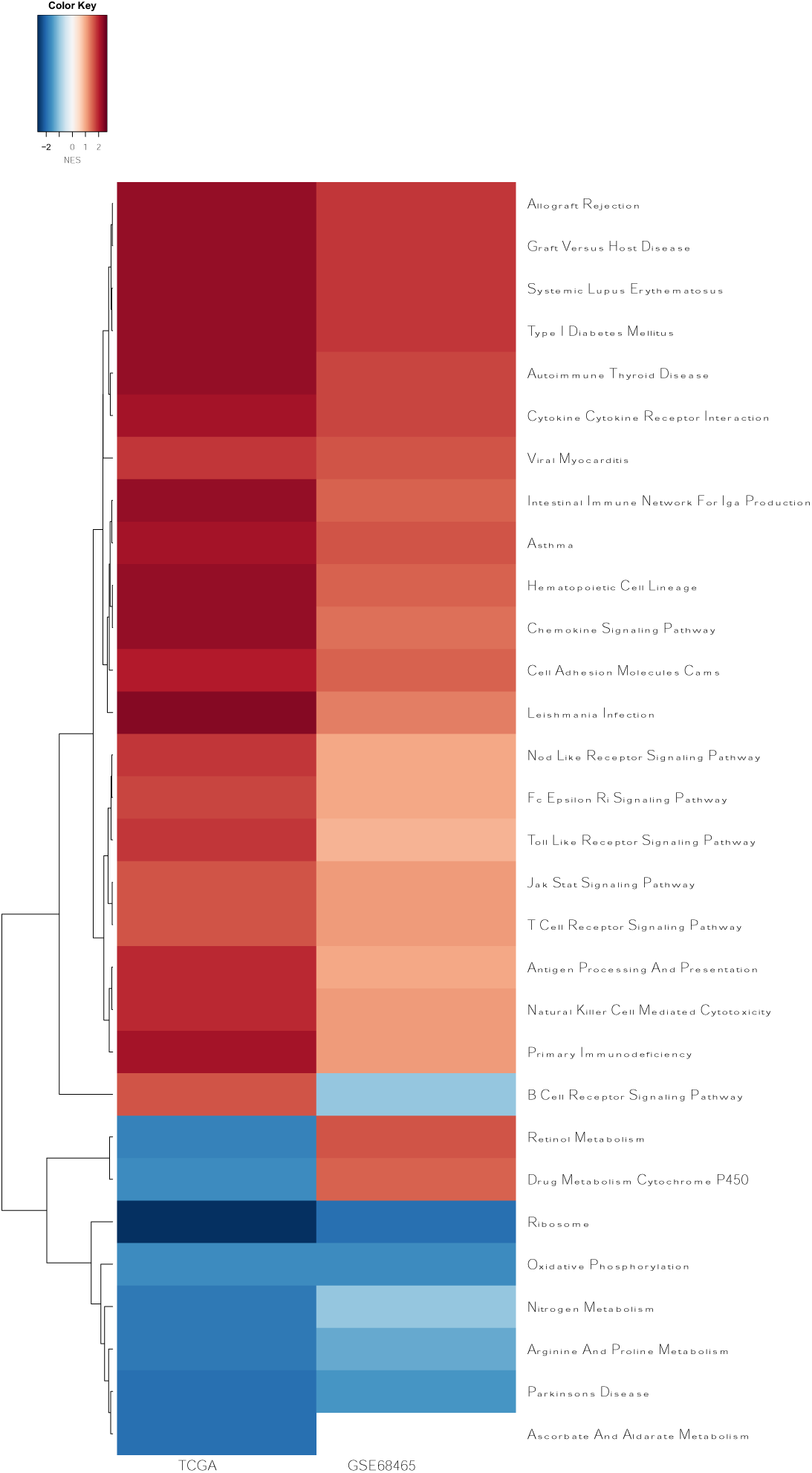
Normalized enrichment scores (NES) of the biological pathways that are significantly (at an FDR cutoff 0.05) differentially targeted with age by transcription factors in tumor samples from TCGA. Left column shows NES from GSEA on TCGA samples, and the right column shows NES for the same pathways from GSEA on GSE68465 samples.

However, there were many pathways with aging-associated regulatory changes found exclusively in tumor samples and not in healthy lung samples, including the NOD-like receptor signaling pathway, FC-epsilon RI signaling pathway, toll-like receptor signaling pathway and JAK-STAT signaling pathway – all of which have been associated with LUAD development, progression and outcome. In contrast, we found that metabolic pathways including oxidative phosphorylation, nitrogen metabolism, arginine and proline metabolism, ascorbate and alderate metabolism were decreasingly targeted by TFs with age. These results were validated in an independent dataset GSE68465 (**Figure 7** right column).

Biological pathways associated with immune response that had been previously identified as increasingly targeted (at an FDR cutoff 0.05) with age in non-cancerous lung samples, also showed increased targeting by TFs with age in tumors. We also found several additional immune-related pathways to have age-dependent regulatory patterns in tumor, that were not evident in non-cancerous samples. Such immune-related pathways include those involved in antigen processing and presentation, graft versus host disease, JAK-STAT signaling pathway, natural killer cell mediated cytotoxicity, primary immunodeficiency and T-cell receptor signaling pathway, all of which were increasingly targeting by TFs with age.

This difference between non-cancerous lung versus LUAD tumor in the age-associated targeting patterns of immune pathways can be partially attributed to differences in immune cell infiltration by age in non-cancerous lung tissue as compared to tumor. Immune infiltration analysis (**Figure C.4**) showed that the proportion of CD8+ central memory cells increased with age in both non-cancerous lungs, and in tumor. However, aging-related changes in immune cell composition in tumor were distinct from those in healthy lungs for most immune cell types. For example, while the proportions of activated myeloid dendritic cells and B cells increased with age in LUAD tumors, in non-cancerous lung the proportion of these cells did not change significantly with age. The proportion of macrophages also increased with age in tumor; in non-cancerous lung, macrophages were more abundant in younger samples. This higher infiltration of immune cells in tumor with age might be associated with a higher targeting of immune pathways among older individuals with LUAD, as evidenced by the positive correlation between immune score and TF-targeting score of immune pathways (**Table C.1**). In contrast, the proportion of CD8+ naïve T-cells and common myeloid progenitors showed a decreasing trend with age in tumor, while exhibiting no significant difference in composition across healthy lung samples of varying age (**Figure C.4**).

### Gene Regulatory Network-informed Aging Signature of Tumor Predicts Survival and Response to Chemotherapy in LUAD

We conducted survival analysis using the Cox proportional hazard model to understand whether the aging-associated regulatory patterns of biological pathways have any influence on the prognosis of LUAD for individuals of varying age. We first constructed an “aging signature” for tumor samples, defined by the outputs of the Cox proportional hazard model with inputs being the targeting score (as defined in Methods) of 28 biological pathways (**Figure 7**) that were discovered to be significantly differentially targeted by TFs as a function of age in tumors. This network-informed “aging signature” of tumors is a linear combination of the TF-targeting scores of 28 biological pathways and is uncorrelated (**Figure C.5**) with both chronological age (correlation = −0.0114 with p-value = 0.792) and clinical tumor stage (p-value from ANOVA = 0.1921).

We found that samples with a lower aging signature had significantly better survival probability than samples with a higher aging signature (**Figure 8** left; p-value = 0.001). For comparison, we split samples into two chronological subgroups based on whether samples were above or below the median chronological age (**Figure 8** right) and not find any significant difference (p-value = 0.169) in survival probability. This indicates that the aging signature, defined using gene regulatory networks is more informative than chronological age in predicting LUAD survival in TCGA. The results remained consistent even after adjusting for self-reported gender, race, smoking status, clinical tumor stage and therapy status (p-values for the network-informed aging signature and chronological age were 2.8e-06 and 0.171 respectively). We validated (**Figure C.6**) the efficacy of the network-informed aging signature in predicting survival outcome in LUAD on independent dataset GSE68465 (p-value = 0.013, after adjusting for gender, race, smoking status, tumor stage and therapy status).

**Figure 8:**
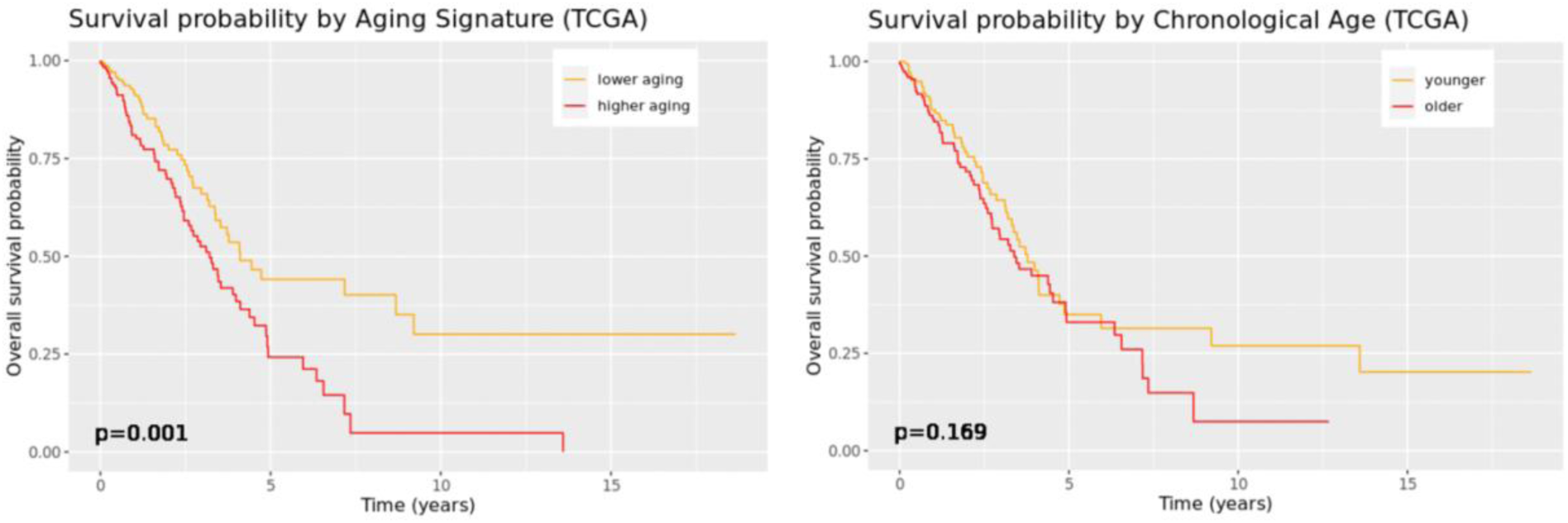
Kaplan-Meier plot for survival outcome in TCGA, split by median network-informed aging signature (left) and by median chronological age into younger and older tumor samples (right).

We also performed a Cox proportional hazard analysis using with therapy information and the aging signature as input (**Table 2**) and found that biological age score captured in the aging signature had significant (p-value = 0.009) interaction with chemotherapy, where a smaller value of the aging signature was associated with higher improvement in survival probability in response to chemotherapy, compared to no therapy. In contrast, using an analogous Cox proportional hazard model with therapy information and chronological age as input, we did not find any significant interaction (p-value = 0.564) between chronological age and chemotherapy response. In the independent dataset GSE68465 (**Table C.2**) as well, the interactions between chemotherapy and the aging signature were in the same direction as in TCGA, although the results (**Table C.2**) were not statistically significant.

### Drug Repurposing with CLUEreg Identifies Distinct Small Molecule Drugs for Tumor Samples with Lower versus Higher Aging Signature

To find potential targeted cancer therapeutics that might differ in efficacy depending on aging signatures, we split the TCGA tumor samples into two groups – one above and one below the median value of the network-informed aging signature. For each group, we separately used CLUEreg [36] to identify small molecule drug candidates depending on the differential regulatory patterns between tumor and healthy samples and obtained a list of 150 small molecule drugs each for the two aging signature groups (**Figure 9**).

**Figure 9:**
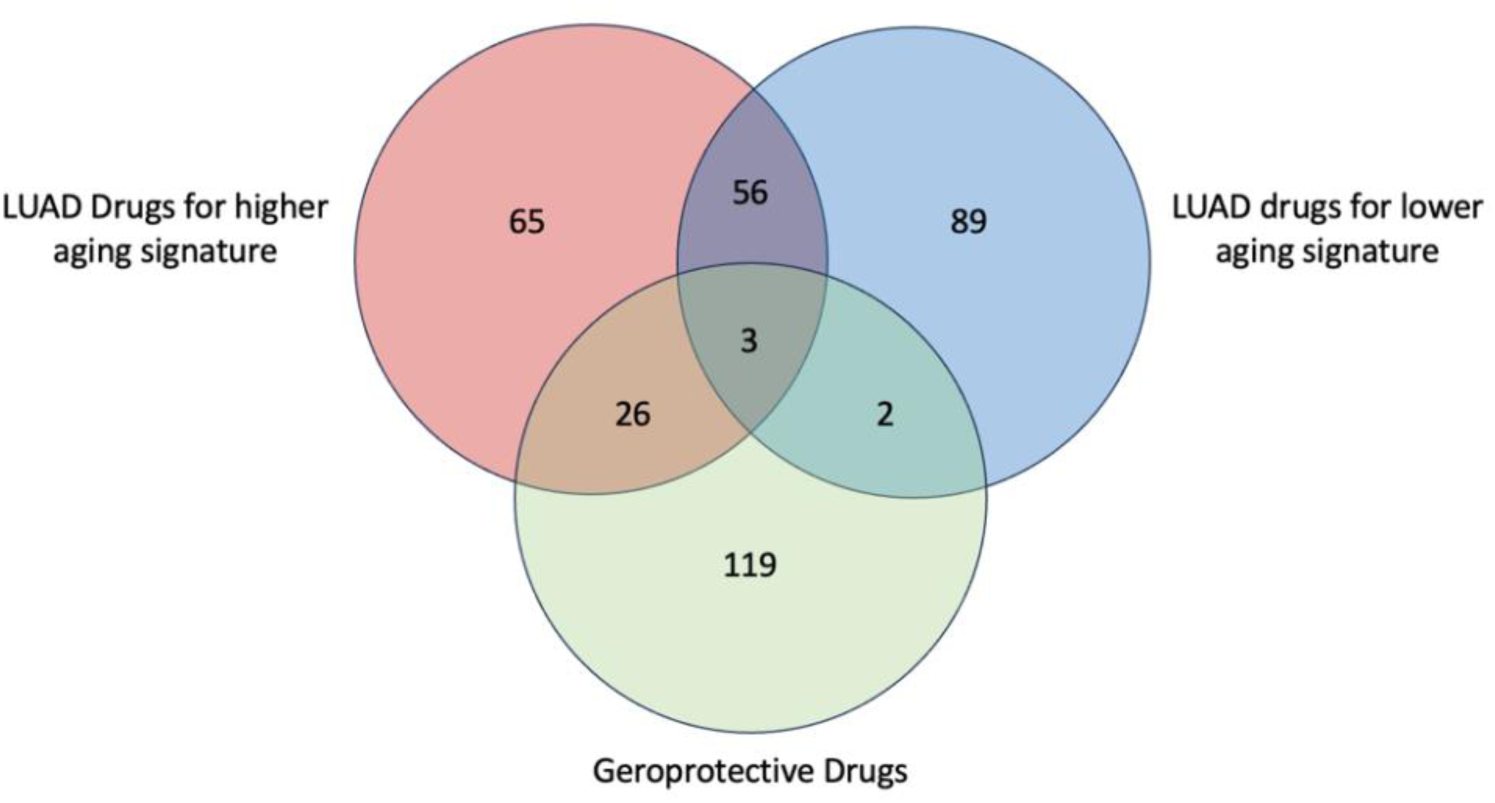
Venn Diagram of Number of Small Molecule Drug Candidates Derived from CLUEreg as (green) geroprotective drugs, (blue) LUAD drugs for individuals with lower aging signature and (red) LUAD drugs for individuals with higher aging signature.

While 59 small-molecule cancer therapeutics including FDA-approved Cisplatin and Amifostine and investigational drugs Timosaponin and Cardamonin appeared as potential drug candidates for both aging-signature groups, several other drugs appeared exclusively in only one of the two groups. Drugs including Homoharringtonine, Ingenol, Vatalanib (investigational), Midostaurin (investigational) and Ubenimex (investigational) appeared only for individuals with lower aging signature. Other drugs including Leucovorin, Actinomycin-d and Plumbagin appeared for the higher aging signature group alone. Several potential geroprotective drug candidates, including Meclofenoxate and Isonicotinamide, also appear in the lists of anti-tumor drugs. It is worth noting that we found 28 geroprotective drugs in the list for higher aging signature group while only 5 geroprotective drugs for the lower aging signature group. This suggests that the information captured by the aging signature encompasses disease-relevant processes driving LUAD, which are intertwined with other aging-related processes that also contribute to the development and progression of the disease.

## Discussion

LUAD, like most other solid tumors, is an age-biased disease in which individuals generally have a greater risk, poorer prognosis and poorer response to most therapies compared to their younger counterparts [4]. However, tumors diagnosed in younger individuals are often detected at more advanced clinical stages implying more severe disease biology. Earlier studies have found that across diverse cancer types, the mutational landscape of tumors in younger individuals is very different from that in older individuals [10]. However, mutational burden alone does not fully explain the mechanisms of disease, driven by the activity of biological pathways that are activated leading up to and during disease. To bridge this gap in our understanding of LUAD, we investigated how regulation of various genes and biological pathways change with age using individual specific gene regulatory networks. Analyzing gene regulatory networks that integrate gene expression, TF-binding motif and TF protein-protein interaction data from both non-cancerous human lung and LUAD samples, we found aging-associated alterations in regulatory mechanisms involving key biological processes.

Analyzing gene regulatory networks in GTEx lung samples, we found several genes with known relevance to cancer incidence and prognosis that were differentially targeted by TFs with age, including proto-oncogenes *AKT1*, *ERBB3* and *MYCN*. Using pathway enrichment analysis, we saw a clearer picture of how age affects cancer-related processes in non-cancerous lung tissue, including altered regulation of intracellular adhesion, cell proliferation, and immune response. By conducting a differential targeting analysis on non-cancerous lung samples and LUAD tumor samples, we confirmed that targeting of these same biological processes also changes in tumor incidence and in the same direction as they do in “normal” aging. This suggests that aging-associated alterations in gene regulatory patterns of these pathways might be a contributing factor to a higher risk of LUAD development among older individuals. Further, we found that tobacco smoking was associated with an acceleration in the aging-associated gene regulatory changes, helping to explain the increased the risk of LUAD incidence at a younger age among individuals with a history of smoking.

Gene regulatory network analysis of LUAD tumor samples identified an age-associated higher targeting of several biological pathways associated with immune response, these associations were not detected in non-cancerous lung samples from GTEx. Greater targeting of immune pathways with age was correlated with a higher infiltration of immune cells including active myeloid progenitors, B cells, macrophages and CD8+ central memory cells. We suspect that a higher targeting of immune pathways in conjunction with a higher proportion of immune cells among older individuals might contribute to an age-biased response to immunotherapy. This is concordant with evidence from earlier studies which demonstrated that while chemotherapy is more beneficial for younger individuals [5], some immune checkpoint inhibitors provide greater benefit to adults with age 65 or older, compared to younger adults [54, 55].

We constructed a network-informed "aging signature" for tumor samples, based on the TF-targeting patterns of 28 biological pathways (identified in TCGA and validated in GSE68465) that exhibited significant differential targeting by TFs with age. We found that individuals with a lower aging signature not only had better survival outcomes, but also had significantly better improvements in survival outcomes in response to chemotherapy compared to those with a higher aging signature. Within TCGA LUAD samples, this network-based aging signature appears to be a possibly superior biomarker to chronological age, in distinguishing between individuals with varying potential for chemotherapeutic efficacy.

The consistent theme of age-associated alterations in regulation being linked to LUAD suggested that network-based aging signatures might identify aging-related tailored therapeutics. Separately analyzing LUAD samples partitioned into low-and-high-aging signature groups, we found 59 small-molecule cancer therapeutics, including FDA-approved Cisplatin and Amifostine, common to both aging-signature groups. But we also found several drugs were exclusive to one aging signature group alone, meaning that considering age-related regulatory changes might be useful in determining personalized therapeutic protocols. Certain potential geroprotective drugs including Meclofenoxate and Isonicotinamide appeared in the lists of anti-tumor drugs, mostly for the higher aging signature group, as did a number of candidate drugs such as Curcumin [38], that have been shown have geroprotective effects. Unfortunately, older adults are severely underrepresented in clinical trials for most cancers including LUAD, thereby impacting the validity of clinical guidelines in diseases with an age effect [56]. Our analysis underscores the importance of including individuals across the spectrum of disease-associated demographics in clinical trials.

It is important to note that our analysis is based on observational data alone and hence experimental validation is required to establish a causal relationship between the aging-associated regulatory changes identified by our analysis and the manifestation of tumor. Another limitation of our study is that the datasets used for discovery (GTEx and TCGA) and validation primarily consist of individuals of white and African American descent. Despite adjusting for the impact of race in our analysis, the applicability of our findings to other ethnicities may still be constrained due to underrepresentation in our data and the confounding effects of various social determinants of health on lung cancer. Further studies involving more diverse populations are necessary to confirm the validity of our results across a broader range of racial and ethnic backgrounds. Additionally, a more complete inclusion of social determinants such as individual socio-economic background [57], is essential for the generalizability of our findings.

Despite these limitations, our analysis provides interesting insights into the aging-associated alterations in gene regulation and their relevance in the clinical manifestation of LUAD, including some immediate implications in the context of personalized cancer therapy. Based on our analysis we infer that aging related changes in regulation of key biological processes involved in intra-cellular adhesion, cell proliferation and immune responses are associated with altered risk, prognosis and response to therapy in lung adenocarcinoma among individuals of varying age. Notably, we observed that even among individuals of similar age, individuals with lower network-informed aging signature had better prognostic outcome in response to chemotherapy, than individuals with higher aging signature. This observation implies that chronological age alone does not provide substantial information on prescribing personalized therapy for lung adenocarcinoma and gene regulatory networks can prove to be effective tools in facilitating more efficient personalized therapy design and improving prognosis in lung adenocarcinoma for individuals across varying age.

What emerges from our analysis is an interesting picture of how ageing influences LUAD. “Normal” aging in the lung is associated with alterations in the regulation of particular biological processes, and indeed, by inferring and analyzing gene regulatory network structure, we identified genes and biological processes that exhibit altered patterns of regulation with age. But as has been known, not all individuals age at the same rate. When we examine LUAD, we find that greater changes in age-associated patterns of gene regulation are more strongly associated with disease than is chronological age. We also find that smoking results in an apparent acceleration of the aging-associated patterns of gene regulation in both the lungs of “healthy individuals with a history of smoking” and in “normal adjacent” tissue from individuals suffering from LUAD, consistent with the fact that smoking dramatically increases risk, progression, and severity, and affects response to therapy. This suggests that the regulatory changes that are captured in the aging signature we derived are, at the least, correlated if not causally linked to LUAD disease processes. Looking at younger people with LUAD, we find that they also exhibit age-acceleration in their “normal” lung tissue relative to their peers without LUAD. Differential regulation in tumors of younger people with LUAD represented a subset of the changes we saw in the tumors of older patients, which suggests that the pathways associated with these changes might be particularly important in understanding the severity of disease in younger individuals. Finally, we found that even among individuals of similar age, those with a lower network-informed aging signature had better prognostic outcome in response to chemotherapy, than individuals with higher aging signature. What all these means is that while chronological aging might have some effect on the risk of developing LUAD and its properties, the changes in aging-related regulation are far more important in estimating disease risk, in understanding disease processes, in identifying candidate therapies, and in designing aging-aware precision treatment protocols. This observation implies that chronological age alone does not provide substantial information on prescribing personalized therapy for lung adenocarcinoma and gene regulatory networks can prove to be effective tools in facilitating more efficient personalized therapy design and improving prognosis in lung adenocarcinoma for individuals across varying age.

## Funding

This work was supported by grants from the National Institutes of Health: **ES, CMLR, MBG, VF, KHS, and JQ** were supported by R35CA220523; **MBG** and **JQ** were also supported by U24CA231846**; JQ** received additional support from P50CA127003; **JQ** and **DLD** were supported by R01HG011393; **KHS** and **DLD** were supported by R01HG125975 and P01HL114501; **DLD** was also supported by K24HL171900; **KHS** was supported by T32HL007427; **CMLR** was supported by K01HL166376; **CMLR** and **ES** were also supported by the American Lung Association grant LCD-821824.

## Author Contributions

**ES:** Conceptualization, Data curation, Formal analysis, Investigation, Methodology, Software, Validation, Visualization, Writing – original draft; **MBG:** Conceptualization, Writing – review & editing; **VF:** Conceptualization, Writing – review & editing; **KHS:** Conceptualization, Writing – review & editing; **DLD:** Conceptualization, Funding acquisition, Supervision, Writing – review & editing; **JQ:** Conceptualization, Funding acquisition, Resources, Supervision, Writing – review & editing; **CMLR:** Conceptualization, Data curation, Funding acquisition, Supervision, Writing – review & editing.

## Declaration of interests

The authors declare no competing interests.

## Data and Code Availability

Raw data to construct gene regulatory networks and other analysis were downloaded from open-source databases dbGap, Recount3, GEO, STRINGdb, CIS-BP and GDSC. Processed data are available upon request.

Sample-specific gene regulatory networks are stored in an Amazon Web Services s3 bucket and will be made available upon acceptance.

R codes for all downstream analysis are available on a GitHub public repository: https://github.com/Enakshi-Saha/Aging-LUAD

A notebook describing differential targeting analysis and computation of network-informed aging signatures using LUAD tumor samples from TCGA will be available on Netbooks [58]: http://netbooks.networkmedicine.org upon acceptance.

## Methods

### Discovery Dataset

Uniformly processed RNA-Seq data were downloaded from the Recount3 database [59] for two discovery datasets using R package “recount3” (version 1.4.0): (i) lung tissue samples from the Genotype Tissue Expression (GTEx) Project [60] (version 8) and (ii) lung adenocarcinoma samples from The Cancer Genome Atlas (TCGA) [61] on May 26, 2022. We accessed clinical data for GTEx samples from the dbGap website (https://dbgap.ncbi.nlm.nih.gov/) under study accession phs000424.v8.p2. Clinical data for TCGA samples were downloaded from Recount3. We refer to the GTEx samples as “non-cancerous lung samples” throughout our analysis.

From 655 lung samples, 71 samples were removed because they were designated as “biological outliers” in the GTEx portal (https://gtexportal.org/) for various reasons (as described in https://gtexportal.org/home/faq). We analyzed the remaining 584 lung samples (187 female and 397 male) from GTEx. We removed two recurrent tumor samples from the TCGA data and included the remaining 541 primary tumor samples (293 female and 248 male) and 59 normal adjacent (34 female and 25 male) samples.

**Table 1** summarizes the clinical characteristics of the datasets.

**Table 1:** Clinical characteristics of the discovery and validation datasets.

|  | GTEX | TCGA | TCGA | LGRC | GSE68465 |
| --- | --- | --- | --- | --- | --- |
| Type | Healthy | Tumor | Normal adjacent | Healthy | Tumor |
| Sample size | 584 | 539 | 59 | 108 | 443 |
| Age |  |  |  |  |  |
| Mean ± std (range) | 54 ± 11.81<br>(21-70) | 65 ± 9.91<br>(33-88) | 66 ± 10.83<br>(42-86) | 64 ± 11.35<br>(32-87) | 64 ± 10.1<br>(33-87) |
| Gender |  |  |  |  |  |
| Female (%) | 187<br>(32.02%) | 291<br>(54.00%) | 34<br>(57.63%) | 59<br>(54.63%) | 220<br>(49.66%) |
| Male (%) | 397<br>(67.98%) | 248<br>(46.00%) | 25<br>(42.37%) | 49<br>(45.37%) | 223<br>(50.34%) |
| Race |  |  |  |  |  |
| White (%) | 499<br>(85.44%) | 411<br>(76.25%) | 55<br>(93.22%) | - | 295<br>(66.60%) |
| Black or African American (%) | 70 (11.99%) | 53 (9.83%) | 4 (6.78%) | - | 12 (2.71%) |
| Others (%) | 15 (2.57%) | 75<br>(13.92%) | 0 | - | 7 (1.58%) |
| Unknown (%) | - | - | - | - | 129<br>(29.11%) |
| Smoking status |  |  |  |  |  |
| History of smoking (%) | 386<br>(66.10%) | 448<br>(83.11%) | 46<br>(77.97%) | 65<br>(60.19%) | 300<br>(67.72%) |
| No history of smoking (%) | 182<br>(31.16%) | 77<br>(14.29%) | 7 (11.86%) | 32<br>(29.63%) | 49 (11.06%) |
| NA (%) | 16 (2.74%) | 14 (2.60%) | 6 (10.17%) | 12<br>(10.18%) | 94 (21.22%) |
| Tumor stage |  |  |  |  |  |
| I (%) | - | 295<br>(54.73%) | 30<br>(50.85%) | - | 150<br>(33.86%) |
| II (%) | - | 126<br>(23.38%) | 13<br>(22.03%) | - | 251<br>(56.66%) |
| III (%) | - | 84<br>(15.59%) | 13<br>(22.03%) | - | 28 (6.32%) |
| IV (%) | - | 26 (4.82%) | 2 (3.39%) | - | 12 (2.71%) |
| NA (%) | - | 8 (1.48%) | 1 (1.69%) | - | 2 (0.45%) |
| **Ischemic time (hours)** |  |  |  |  |  |
| Mean ± std (range) | 8.04 ± 6.98<br>(0.0-24.4) | - | - | - | - |

**Table 2:** Cox proportional hazard model in TCGA to predict survival outcome using therapy status and network-informed aging signature.

| Covariate | Coefficient | z-score | P-value |
| --- | --- | --- | --- |
| Aging_signature | 1.330 | 6.093 | 1.1e-09 |
| TherapyChemotherapy | 0.445 | 2.059 | 0.040 |
| TherapyOther | -0.279 | -0.277 | 0.782 |
| Aging_signature: TherapyChemotherapy | **-0.905** | **-2.623** | **0.009** |
| Aging_signature: TherapyOther | -1.338 | -0.615 | 0.539 |

Both GTEx and TCGA gene expression data were normalized by transcript per million (TPM), using the “getTPM” function in the Bioconductor package “recount” (version 1.20.0) [62] on R version 4.1.2. Lowly expressed genes were filtered out by removing genes with counts <1 TPM in at least 10% of the samples (126 samples) in GTEx and TCGA combined, removing 36386 genes, and keeping 27470 genes. To construct gene regulatory networks, we further removed those genes that were not present in the TF/target gene regulatory prior used for creating the gene regulatory networks (see section “Differential targeting analysis using sample-specific gene regulatory networks”). This filtering left with 27162 genes, including those on allosomes, which were used in subsequent analysis. For female samples in both GTEx and TCGA, gene expression values of all genes on the Y chromosome (36 genes in total) were replaced by “NA”. Principal component analysis did not show any visible batch effect.

### Validation Dataset

We chose two independent studies for validation from the Gene Expression Omnibus (GEO) repository: GSE47460 [63] (downloaded on Feb 12, 2023) and GSE68465 [64] (downloaded on Jan 24, 2023). For validating our results on the lung samples from GTEx, we used GSE47460, which consisted of microarray gene expression for 582 samples in total from the Lung Genomics Research Consortium (LGRC). This study used Agilent-014850 Whole Human Genome Microarray 4×44K G4112F and Agilent-028004 SurePrint G3 Human GE 8×60K Microarray for sequencing. Among these 582 samples, we used only 108 samples (59 female and 49 male), who have no chronic lung disease by CT or pathology and hence, were designated as “controls” within the study. GSE68465 consists of microarray gene expression for 443 lung adenocarcinoma samples (220 female and 222 male). This study used Affymetrix Human Genome U133A Array for sequencing.

**Table 1** summarizes the clinical characteristics of the datasets analyzed.

Normalized expression data and clinical data were downloaded from GEO using R package “GEOquery” version 2.62.2. Within every dataset, for genes with multiple probe sets, we kept the probe set with the highest standard deviation in expression levels across samples. We discarded any genes that were not in the TF/target gene regulatory network prior that we used for creating the gene regulatory networks. This process left 13575 genes in GSE47460 (LGRC) dataset and 11725 genes in GSE68465 dataset, that we used to build gene regulatory network models. The LGRC data were not batch corrected because no visible batch effect was detected from principal component analysis. The GSE68465 dataset includes lung adenocarcinoma specimens from the following sources: University of Michigan Cancer Center (100 samples), University of Minnesota VA/CALGB (77 samples), Moffitt Cancer Center (79 samples), Memorial Sloan-Kettering Cancer Center (104 samples), and Toronto/Dana-Farber Cancer Institute (82 samples). The GSE68465 data were batch corrected for these sources using “ComBat” function implemented in the R package “sva” (version 3.42.0).

### Differential Targeting Analysis using Sample-specific Gene Regulatory Networks

Gene regulatory networks for each sample were reconstructed by the PANDA [19] and LIONESS [20] algorithms using Python package netZooPy [65] version 0.9.10, in both the discovery and the validation datasets. A schematic diagram of our network construction pipeline is given in**Figure 1**. Three types of data were integrated to derive the regulatory networks: TF/target gene regulatory prior (obtained by mapping TF motifs from the Catalog of Inferred Sequence Binding Preferences (CIS-BP) [66] to the promoter of their putative target genes), TF protein-protein interaction data (using the interaction scores from StringDb v11.5 [67] between all TFs in the regulatory prior), and gene expression (from the discovery or validation datasets). The TF/target gene regulatory prior contains regulatory edges from 997 TFs to 61485 ensemble gene IDs, corresponding to 39618 gene symbols (HGNC, Gencode v39). The protein-protein interaction data contained measures of interactions between the 997 TFs. We used sex-specific motif priors for male and female samples, where the motifs coincided for autosomal genes and genes on the X chromosome and differed for genes on the Y chromosome. The procedure for deriving the motif prior and the PPI priors are described in the Supplementary Material.

Regulatory networks were constructed independently for each of the discovery and validation datasets, and separately for female and male samples. The final networks contained only genes overlapping between the TF/target gene motif prior and the corresponding gene expression dataset.

For each sample’s gene regulatory network, we calculated the targeting score for each gene, equivalent to the gene’s in-degree (defined as the sum of all incoming edge weights from all TFs in the network). The resulting gene targeting scores were compared across individuals of varying ages, using a linear regression model, while correcting for relevant covariates, using the R package limma (version 3.50.3) [51]. The resulting t-statistics of the age coefficient are then used for a gene set enrichment analysis (GSEA).

#### Model 1

To investigate how aging influences the targeting of different genes in the non-cancerous lung samples, we fit a linear model using R package “limma” using the gene targeting score of all genes in GTEx as response and age as covariate, while adjusting for self-reported gender (Male and Female), race (White, Black or African American and others), smoking status (individuals who have never smoked in their lifetime and individuals with a current or past history of smoking), RNA integrity number (RIN), batch and ischemic time. A similar analysis was replicated in the LGRC validation data where we adjusted for gender and smoking status.

#### Model 2

In a separate analysis, to identify biological processes relevant for cancer development, we compared the gene regulatory networks constructed with non-cancerous lung samples from GTEx to the networks constructed from LUAD tumor samples from TCGA with another linear model fit by “limma” using the following covariates: age (“age at diagnosis”), disease status (healthy vs tumor), self-reported gender (Male and Female), race (White, Black or African American and others) and smoking status (individuals who have never smoked in their lifetime and individuals with a current or past history of smoking). The resulting t-statistics corresponding to disease status quantifies the difference between TF-targeting in tumor versus healthy samples and were subsequently used for gene set enrichment analysis (GSEA).

#### Model 3

To identify which biological processes are differentially regulated with age among LUAD tumor samples, we fit a linear model on the indegree of genes computed from the GRNs constructed on LUAD tumor samples from TCGA and derive the t-statistics of the regression coefficients corresponding to age (at diagnosis of LUAD), while controlling for clinical variables such as self-reported gender, race, smoking status and tumor stage. A gene set enrichment analysis was performed using the ranked t-statistics of the age coefficients derived from the limma analysis.

### Pathway Enrichment Analysis

We performed pathway enrichment analysis (**Figure 1**) with a pre-ranked Gene Set Enrichment Analysis (GSEA) using R package “fgsea” (version 1.20.0) [68] and the gene sets obtained from the Kyoto Encyclopedia of Genes and Genomes (KEGG) pathway database [69] (“c2.cp.kegg.v2022.1.Hs.symbols.gmt”) that were downloaded from the Molecular Signatures Database (MSigDB) (http://www.broadinstitute.org/gsea/msigdb/collections.jsp). After filtering out those genes not in the expression dataset, only gene sets of size greater than 15 and less than 500 were considered; this restricted our analysis to 176 gene sets. All genes were ranked by the t-statistic produced by the “limma” (version 3.50.3) differential targeting analysis after adjusting for covariates. The resulting ranked set (with gene symbols corresponding to Gencode v39) was used as input to the GSEA. We performed multiple testing corrections using the Benjamini-Hochberg procedure [70].

### Constructing Gene Set and Smoking History-specific Aging Trajectories for Non-cancerous Samples

From our analysis on non-cancerous lung samples from GTEx, we identified genes of two categories: genes that are increasingly targeted by TFs with age (identified by positive age coefficients in the limma model) and the remaining genes that are decreasingly targeted by TFs with age (identified by negative age coefficients in the limma model). Based on these two gene sets, we construct two aging trajectories.

For each of these sets of genes, an “aging trajectory” was constructed: for every sample we computed the mean indegree of all the genes in the set and designated this number as the TF-targeting score for that sample. Then we stratified all the samples in a particular dataset into 20 consecutive age-groups so that each group contained 5% of all samples in the data. Within each age group, we divided all individuals into subgroups based on their smoking history and computed the medians of the gene targeting scores across individuals in each of the smoking subgroup. For each set of genes, a scatterplot (**Figure C.1**) of the gene targeting scores (y-axis) over the mid-point of each age-group (x-axis) was drawn and colored by smoking history. The scatterplots also show the linear regression lines computed for individuals in different smoking categories. These lines are referred to as the aging trajectories of that group.

### Immune Infiltration Analysis

Tumor immune deconvolution analysis was performed on the TCGA data to investigate how immune cell composition varies in tumor samples over different ages. We used “xcell” [71] on the unfiltered TCGA gene expression data with R package “immunedeconv” (version 2.1.0) to infer immune and stromal cell composition in tumor samples. For every cell type, we fit a linear model to predict the corresponding cell proportion on age, while adjusting for other clinical covariates such as gender, race, smoking status, and clinical tumor stage, thus allowing us to quantify how cell proportion in tumor changes with age. To quantify the relation between the proportion of each cell type and TF-targeting score of immune pathways, we used Pearson’s correlation coefficient between the two quantities and test for significance.

### Constructing a Network-based Aging Signature for Tumor Samples

We constructed a network-based aging signature based on the tumor samples from TCGA as follows. For every biological pathway that was significantly differentially targeted by TFs with age in TCGA dataset, we defined a pathway targeting score as follows. A principal component analysis (PCA) was performed on the indegree of all genes in a particular pathway and the first principal component was defined as an aggregated TF-targeting score for the corresponding pathway. To construct an aging signature for tumor samples, we fit a Cox proportional hazard model to predict survival probability using pathway targeting scores for all pathways found to be differentially targeted by age in both the TCGA and GSE68465 validation dataset. The resulting prediction from the Cox proportional hazard model was defined to be the aging signature for the corresponding tumor sample.

Survival analysis with Cox proportional hazard model was implemented using the R package “survival” (version 3.2.13). We also performed Kaplan–Meier survival analysis as implemented in the same R package, and the p-values were computed using the log-rank test. The survival curves were plotted using the R function “ggsurv” on the “GGally” package 2.1.2.

### Small Molecule Drug Repurposing with CLUEreg

We used a web-based drug repurposing tool CLUEreg [36] (https://grand.networkmedicine.org/), designed to match disease states to potential small molecule therapeutics, based on comparing of input regulatory networks to networks computed using PANDA and LIONESS using data from drug-response assays.

We used linear models (R package “limma”) on gene targeting scores from GTEx to identify genes that were significantly (p-value < 0.05) either increasingly (1018 genes) or decreasingly (404 genes) targeted with age. These differentially targeted genes were used as input to CLUEreg web application. CLUEreg produced a list of 150 small molecule drug candidates most suitable for reversing the alterations in gene targeting patterns associated with aging, which we subsequently refer to as drugs with potential geroprotective (i.e. anti-aging) effects.

We also used CLUEreg to identify small molecule drugs for reversing the gene regulatory patterns in tumor samples into regulatory patterns akin to those in normal samples and identified two separate lists of 150 small molecule drugs each, for tumor samples with lower versus higher aging signatures. To identify genes differentially targeted between tumor and normal adjacent samples in TCGA, we fit linear models (“limma”) on gene targeting scores on sample type (tumor versus normal adjacent), while adjusting for other clinical covariates age, gender, race, smoking status and clinical tumor stage. We categorized all samples into lower versus higher aging signature groups by splitting them into two equal parts by median aging signature. We included an interaction term between sample type and aging signature group in the limma analysis to capture the tumor-associated gene regulatory changes in the two aging signature groups. The resulting list of significantly (p-value < 0.05) differentially targeted genes between tumor and normal adjacent samples were used as input to CLUEreg.

## Supplementary Material

### A. Designing Sex-specific Transcription Factor-Gene Motif Prior

The prior regulatory network used in PANDA is a bipartite network consisting of transcription factors and their target genes, where the edges (0 or 1) indicate whether a transcription factor motif exists in a target gene’s promoter region. To create the prior regulatory network, we downloaded Homo sapiens transcription factor motifs with direct/inferred evidence from the Catalog of Inferred Sequence Binding Preferences CIS-BP Build 2.0 (http://cisbp.ccbr.utoronto.ca). We mapped these transcription factor position weight matrices (PWM) to the human genome (hg38) using FIMO [72] and retained highly significant matches (p<10-5) that occurred within the promoter regions of Ensembl genes (Gencode v39; annotations downloaded from http://genome.ucsc.edu/cgi-bin/hgTables); promoter regions were defined as [-750; +250] base pairs around the transcription start site (TSS). This process produced an initial map of potential regulatory interactions involving 997 transcription factors targeting 61,485 genes. To statistically compare networks, the same set of edge combinations need to be included in both sexes, therefore we created sex-informed transcription factor regulatory priors to account for the lack of Y chromosome genes in females. In the female regulatory prior, edges from or to Y chromosome genes were downweighed to zero, which consisted of 52,266 edges.

### B. Designing Protein-protein Interaction Prior

We used the STRINGdb Bioconductor package [73] to access and download PPI data from the StringDB database (STRING.version 11.5). We filtered the PPI data to keep only those interactions present between transcription factors in the prior network (score threshold index of 0). PPI scores were normalized by dividing them by 1000 to have a uniform range between 0 and 1 for the PPI and the prior network. We set transcription factor self-interaction equal to one for all 997 TFs. Since PPI are undirected, we converted the data into a symmetric form.

### C. Additional Tables and Figures

**Table C.1:**
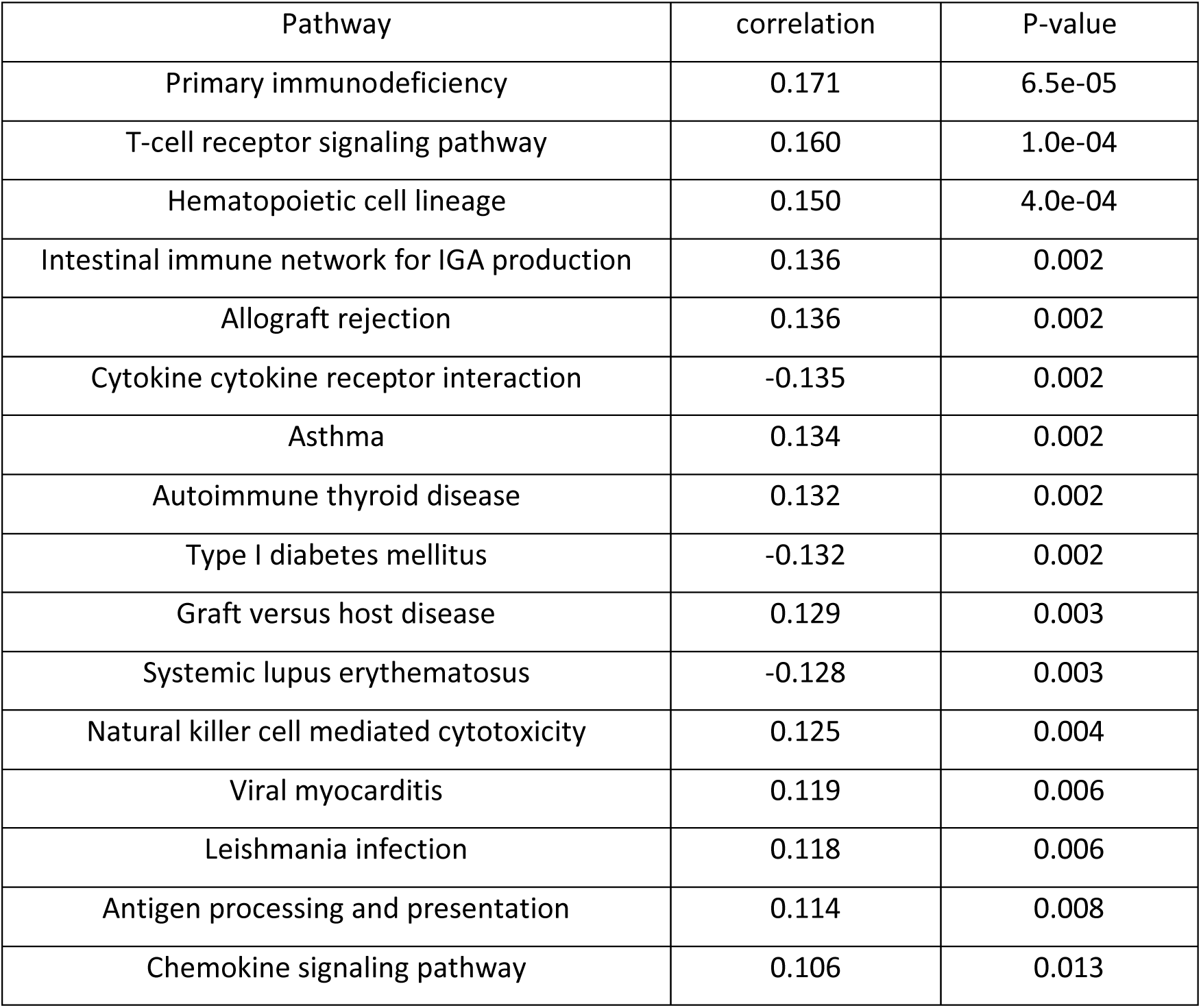
Correlation between pathway targeting score of immune pathways and immune score computed by “xcell”. Only significant correlations (p-value < 0.05) are reported.

**Table C.2:** Cox proportional hazard model in GSE68465 to predict survival outcome using therapy status and network-informed aging signature.

| Covariate | Coefficient | z-score | P-value |
| --- | --- | --- | --- |
| Aging_signature | 0.119 | 2.375 | 0.018 |
| TherapyChemotherapy | 0.467 | 3.123 | 0.002 |
| TherapyUnknown | 0.369 | 1.074 | 0.283 |
| Aging_signature: TherapyChemotherapy | **-0.037** | **-0.516** | **0.606** |
| Aging_signature: TherapyOther | -0.289 | -1.906 | 0.057 |

**Figure C.1:**
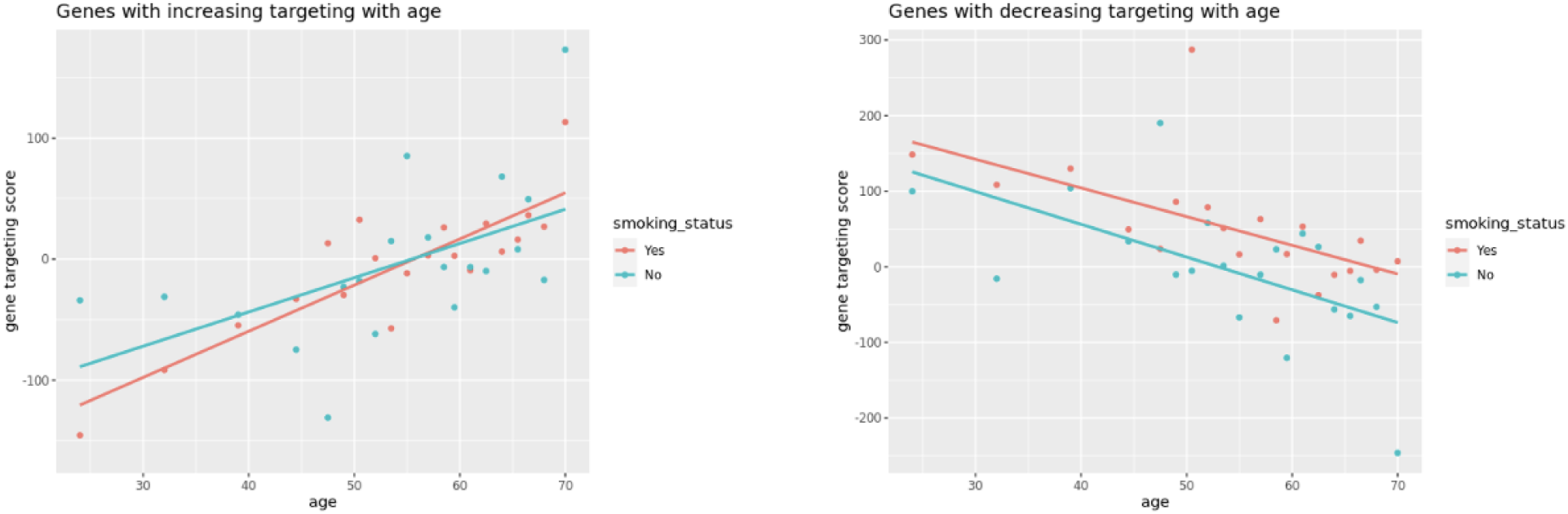
Aging trajectories for the GTEx samples based on two sets of genes. The plot on the left shows an aging trajectory for smokers (current and past smokers with smoking status = “yes”) and lifelong nonsmokers (smoking status = “No”) constructed based on 1018 genes that are significantly increasingly targeted with age by transcription factors. The plot on the right shows an aging trajectory for smokers (current and past smokers with smoking status = “yes”) and lifelong nonsmokers (smoking status = “No”) constructed based on 404 genes that are significantly decreasingly targeted with age.

**Figure C.2:**
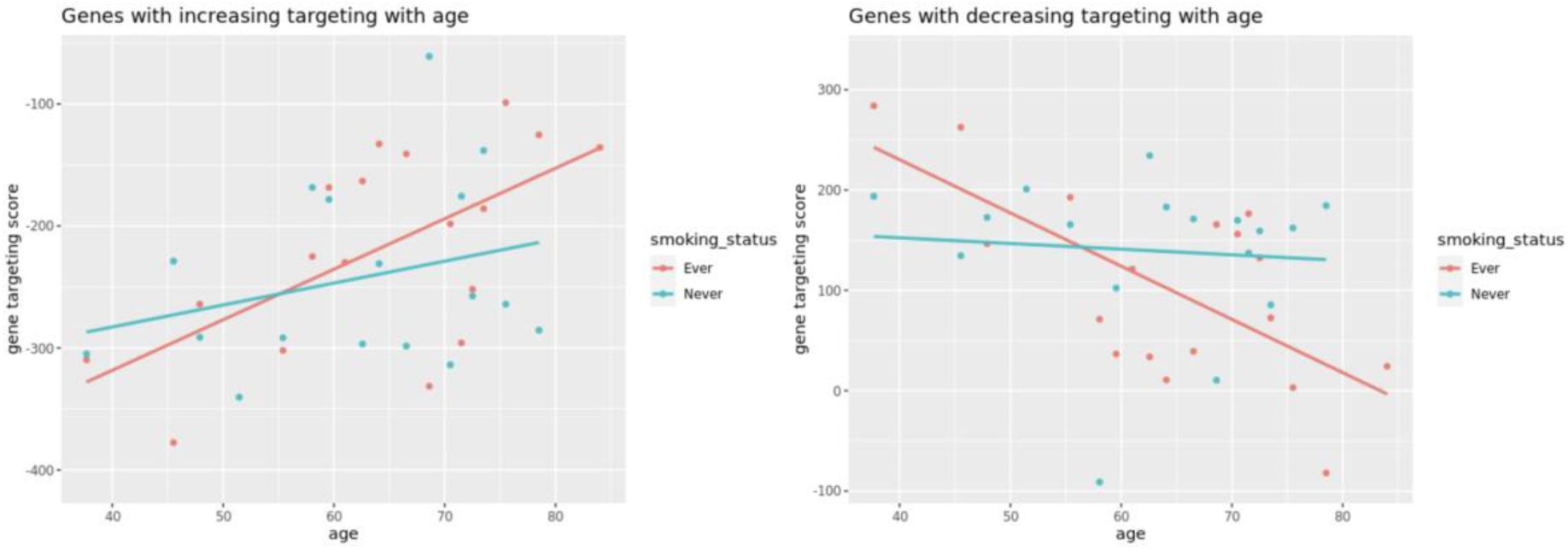
Aging trajectories for the LGRC samples based on two sets of genes. The plot on the left shows an aging trajectory for smokers (current and past smokers with smoking status = “Ever”) and lifelong nonsmokers (smoking status = “Never”) constructed based on 888 genes that are significantly increasingly targeted with age by transcription factors. The plot on the right shows an aging trajectory for smokers (current and past smokers with smoking status = “Ever”) and lifelong nonsmokers (smoking status = “Never”) constructed based on 556 genes that are significantly decreasingly targeted with age.

**Figure C.3:**
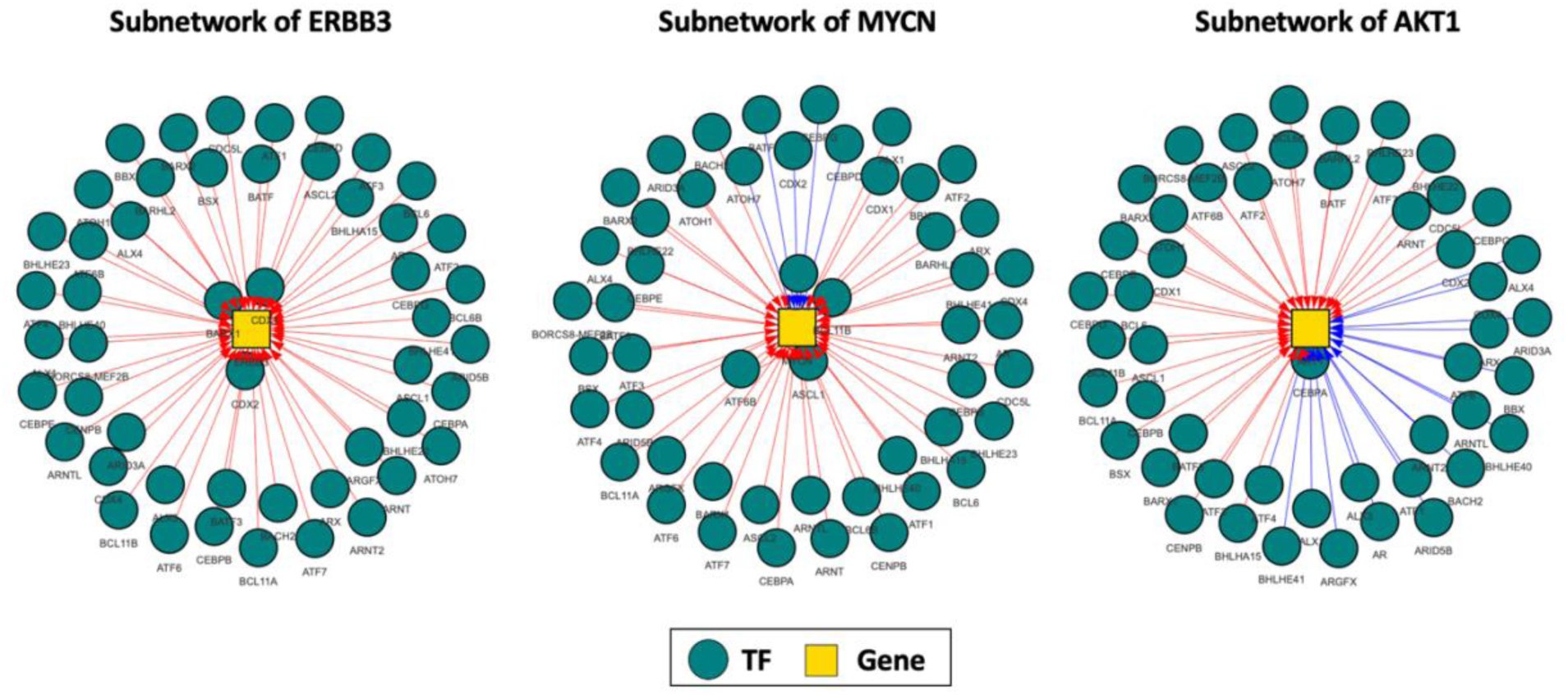
Aging-associated change in TF-targeting patterns of oncogenes ERBB3, MYCN and AKT1 in GTEx. Weights of edges marked in red increase with age and weights of edges marked in blue decrease with age. For each gene top 50 TFs are shown for which the targeting pattern changes most with age.

**Figure C.4:**
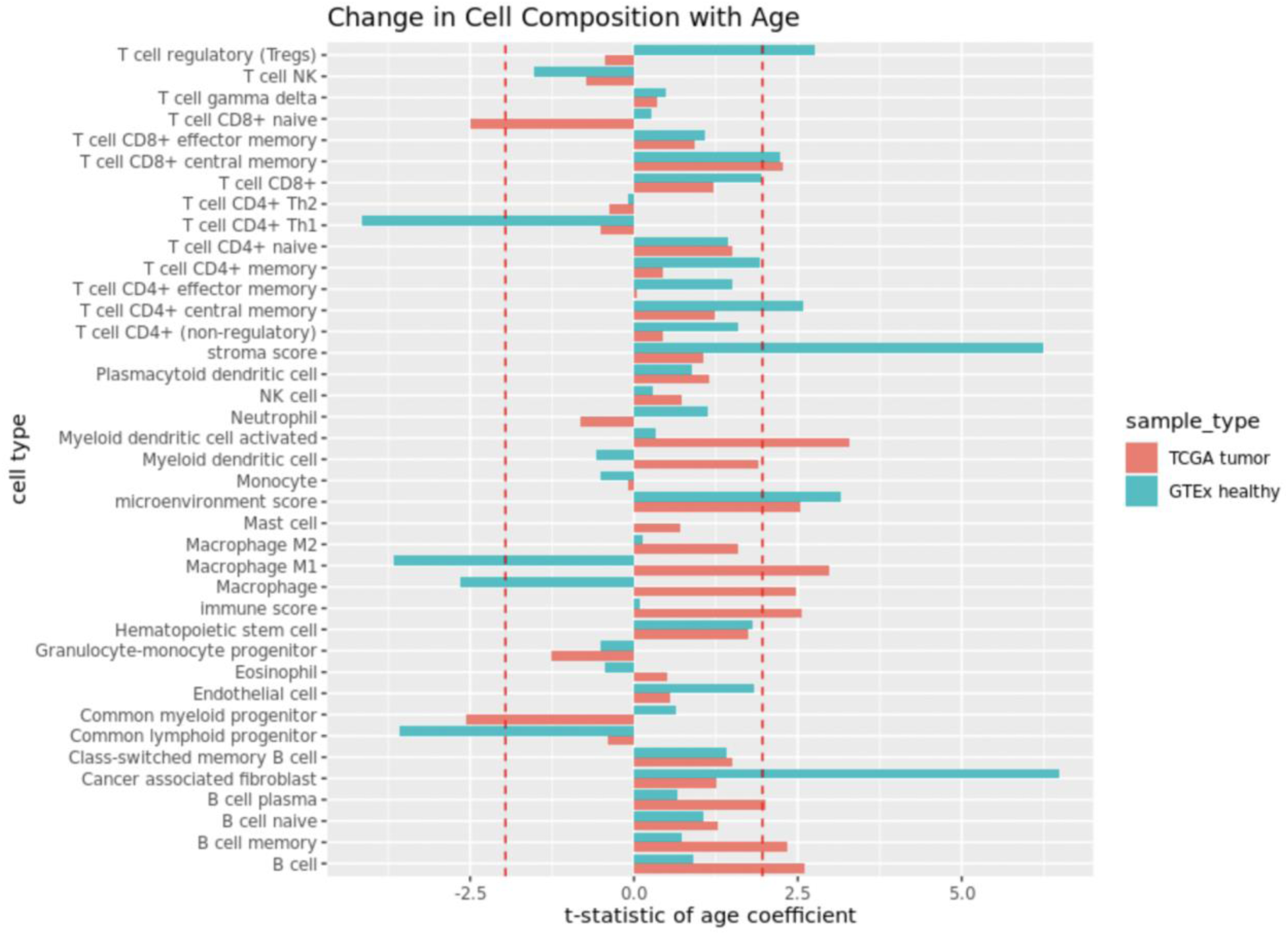
Change in immune and stromal cell composition with age in GTEx and TCGA: for each cell type, the bar lengths correspond to the t-statistics of the age coefficients from linear models with cell type proportion as response and age as covariate, while adjusting for other clinical covariates. Vertical red dotted lines show the 2.5% and 97.5% quantiles of the standard normal distribution. Cell types for which the corresponding bars cross these lines are inferred to be significantly (p-value < 0.05) changing in composition with age.

**Figure C.5:**
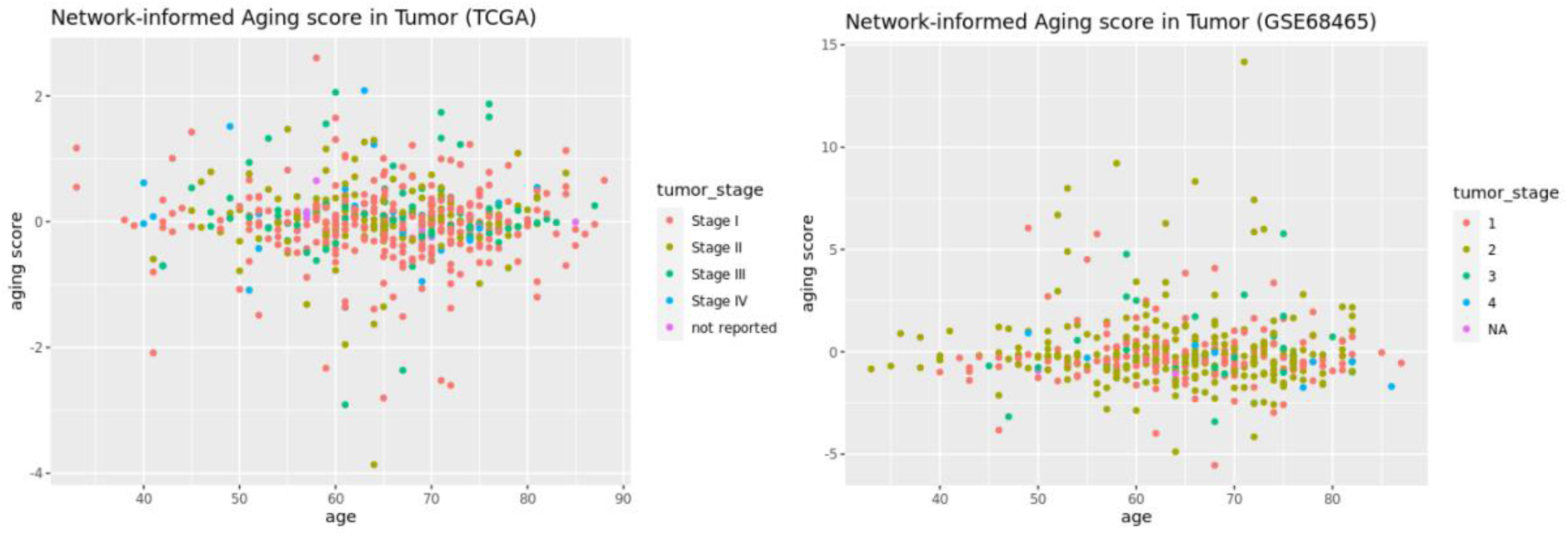
Network-informed aging score versus chronological age, in tumor samples from TCGA (left) and GSE68465 (right), colored by clinical tumor stage.

**Figure C.6:**
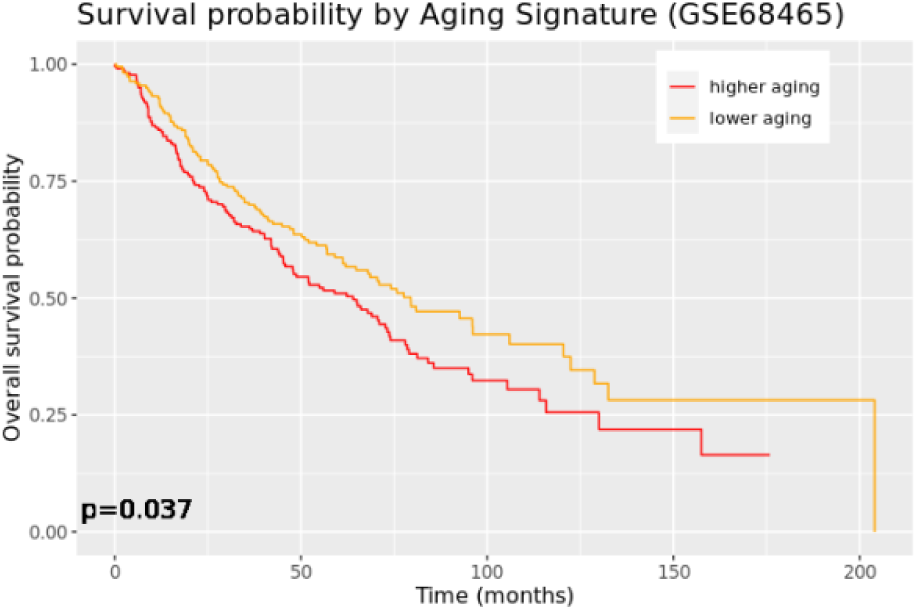
Kaplan-Meier plot for survival outcome in GSE68465, split by lower and higher network-informed aging signature.

## Notes

### Competing Interest Statement

The authors have declared no competing interest.

